# Influence of geography, seasonality and experimental selection on *Chironomus riparius* recombination rates

**DOI:** 10.1101/2025.06.04.657814

**Authors:** María Esther Nieto-Blázquez, Cosima Caliendo, Laura C. Pettrich, Ann-Marie Waldvogel, Markus Pfenninger

## Abstract

**Background:** Understanding recombination rates is crucial in evolutionary biology, as recombination shapes genetic diversity, natural selection, and adaptation. We investigated recombination rate variation in *Chironomus riparius* across different latitudes, seasons, and experimental treatments using Pool-seq data from five studies and the ReLERNN neural network-based method. We examined its relationship with genetic diversity, GC content, and F_ST_, assessing causality through structural equation modeling.

**Results:** In natural populations, recombination rates showed no clear latitudinal pattern, likely due to interactions between climate-driven selection and regional environmental heterogeneity. However, seasonal variation was evident, with higher recombination rates in autumn than winter, possibly due to temperature-induced plasticity or seasonal bottlenecks. A cold snap in March 2018 triggered a sharp recombination increase, potentially suggesting a stress-induced adaptive response. In experimental populations, thermal regimes had no direct effect on recombination, but adaptation to lab conditions was significant. Environmental stressors produced distinct responses: microplastic exposure reduced recombination genome-wide, likely due to stress-induced DNA repair prioritizing genome integrity, while cadmium exposure generally suppressed recombination.

**Conclusions:** Our findings reveal recombination as a highly dynamic process influenced by environment, selection, and genetic background, underscoring the importance of the context in shaping genomic architecture under both natural and experimental conditions.

## BACKGROUND

Life’s diversity emerges from the complex interplay between the environment, evolutionary forces and the genomes they shape. Recombination, the physical exchange of genetic material between homologous chromosomes, is a fundamental evolutionary mechanism that generates genetic diversity, facilitates adaptation, and influences genome evolution [1]. By reshuffling alleles and producing novel genetic combinations, recombination modulates natural selection [2, 3] and contributes to the distribution of genetic variation across the genome [4, 5]. Despite being a conserved meiotic process, recombination rates exhibit substantial variation across genomes, species, and populations due to evolutionary, environmental, and stochastic factors [6].

Recombination rate variation is observed at multiple levels. Closely related species display different recombination landscapes due to distinct genome architectures and evolutionary histories [7, 8]. Within species, recombination rates can vary based on genetic background, environmental conditions, and demographic history of the constituent populations [9–11]. Within populations, recombination rates are influenced by both genetic and stochastic factors, including chromosomal variation [12, 13] and sex-specific differences, with females typically exhibiting higher crossover rates than males [14, 15]. Furthermore, recombination is not uniformly distributed along chromosomes; certain genomic regions, known as recombination hotspots, experience higher crossover frequencies. While recombination does not directly impact an individual’s survival, it shapes genetic variation in offspring and influences the evolutionary trajectory of populations, potentially playing an adaptive role [6, 16–19] - in the same way that variation in fitness-related traits does

A growing body of research highlights the role of biological and environmental factors in driving recombination rate variation. Geographic heterogeneity and latitudinal gradients can shape recombination landscapes by exposing populations to distinct selective pressures and genetic drift regimes [18, 20, 21]. For instance, studies in *Drosophila melanogaster* have revealed regional differences in recombination rates, with local adaptation and demographic history contributing to these patterns [22]. Environmental gradients and ecological clines can also influence recombination, as organisms in fluctuating environments may exhibit elevated recombination rates to enhance adaptive potential [23–25].

Among environmental factors, temperature is a well-documented extrinsic modulator of recombination rates [26–29]. Experimental studies in *D. melanogaster* demonstrate the plasticity of recombination, with increased crossover rates observed in response to temperature fluctuations [30–32]. Beyond temperature, seasonal changes have been shown to impact recombination rates as organisms respond to cyclic environmental stressors. In *D. melanogaster*, crossover rates differ between winter and autumn populations, suggesting that recombination may be influenced by seasonal selection pressures [33]. Theoretical models further suggest that recombination rate modifiers are more likely to persist in fluctuating environments when linked to loci under selection [34]. A substantial body of research has employed experimental setups using non-temperature stressors to investigate recombination, offering critical insights into the mechanisms and factors influencing this fundamental evolutionary process [35–37].

In addition to environmental influences, recombination is correlated with specific genomic features. In honeybees, DNA structure significantly influences recombination frequency, though the mechanisms underlying high recombination rates remain incompletely understood [38]. GC content has been linked to recombination in multiple taxa, including nematodes, *Drosophila*, chickens, and mammals (nematodes and *Drosophila* [39, 40]; *D. melonogaster* [41]; chickens [42]; mouse [43]), though studies in yeast suggest that GC content alone may not always drive recombination [44] or GC content acting as a sole modifier of recombination [45]. Recombination also plays a key role in shaping linkage disequilibrium and nucleotide diversity (π) [3]. Strong correlations between recombination rates and genetic diversity have been documented in *Drosophila* [22, 46–50] and other taxa (butterflies [51]; mice and rabbits [52]; chickens [53]; yeast [54]), though this relationship appears weaker or absent in some plant species [55] and in *D. pseudoobscura* [56].

Another genomic feature associated with recombination is the distribution of transposable elements (TEs; [57–59]. TEs are not randomly dispersed across genomes, and their abundance correlates negatively with recombination rates in *Drosophila*, likely due to stronger purifying selection against TE insertions in high-recombination regions [60–64]. However, this pattern is not universal; in *Caenorhabditis elegans*, recombination rates correlate positively with TE abundance, suggesting lineage-specific variation in the forces governing TE dynamics (RNA-based elements; [65]. Differences in population sizes, selection pressures, and TE families likely contribute to these contrasting patterns [66].

While much of our understanding of recombination comes from model organisms such as *D. melanogaster*, recombination rate variation remains largely unexplored in many non-model species. Furthermore, most studies have focused on spatial variation in recombination, whereas fewer have examined how recombination varies across temporal scales, particularly in response to seasonal or multi-generational environmental changes. Experimental evolution approaches provide a powerful framework to investigate these temporal dynamics, offering insights into how recombination rates evolve under controlled selective pressures. In this study, we conduct an exploratory analysis to investigate recombination rate variation in *Chironomus riparius* using multiple Pool-Seq datasets. Specifically, we aim to examine how recombination rates fluctuate across geographic locations, seasonal changes, and experimental selection regimes. By integrating these datasets, we seek to provide new insights into the forces shaping recombination rate variation and to expand our understanding of the evolutionary and ecological significance of recombination.

## MATERIAL AND METHODS

### Datasets trimming and re-mapping

We download Pool-seq data from ENA for 5 published studies investigating different aspects of the evolutionary and molecular ecology of *C. riparius* (Table S1). *C. riparius* is a multivoltine species of non-biting midge that can produce up to 15 generations annually [67] and is widely distributed in the temperate northern hemisphere [68]. Detailed information on the biology of the species, samples collection experimental setup, and raw Pool-seq data pre-processing can be found in the specific publications (Table S1). Two datasets represent natural populations (geography and seasonality), while three datasets derive from experimental populations (temperature selection, microplastics exposure, and cadmium treatment). Identifiers (IDs) for the seasonality, temperature, microplastics, and cadmium datasets are provided in Table S2. Among the datasets used in this study, only the dataset from [69] was trimmed. For the remaining four datasets, we trimmed adapters and low-quality bases from paired reads with Trimmomatic v.0.39 [70] using 8 threads and a phred score of 33. We assessed quality of reads with using FastQC [71]. Clean reads from all five datasets were mapped to the chromosome level reference genome of *C. riparius* [72] using the BWA mem algorithm [73]. We used samtools v.1.20 *mpileup* module to combine pools for each dataset independently. We used Popoolation2 [74] v.1.201 script *mpileup2sync.pl* to convert each mpileup file into sync files using a minimum quality of 20. Using the functions *read.sync* and *af* from the R package *poolSeq* [75] we calculated allele frequencies (AF) for each pool keeping only for biallelic loci.

### Recombination rates analyses

In order to estimate recombination rates across the genome we used ReLERNN v.1.0.0 [76], a recurrent neural networks (RNN)-based method for estimating the genomic map of recombination rates per chromosome directly from AF for each pool independently. This method provides a more flexible, efficient, and noise-tolerant (derived from uneven coverage and sequencing errors) alternative to traditional methods, especially for Pool-seq data, where phasing is principally not available. Its machine learning approach allows it to adapt better to diverse evolutionary scenarios. First, we used ReLERNN_SIMULATE to split the input of AF and ran the simulations using a window size of 10kb, which included simulate training, validation, and test using coalescent simulation program *msprime*. In a second step, ReLERNN_TRAIN used the simulations from the previous step to train a recurrent neural network. We used 250 epochs to train over and 10 validation steps. Following the training, we used the module ReLERNN_PREDICT to predict per-base recombination rates in 10kb non-overlapping windows across all chromosomes of the genome. Finally, we obtained 95% CI (confidence interval) around each predicted recombination rate with the module ReLERNN_BSCORRECT. For each dataset, we inspected if recombination rates were statistically different with the R package *Bayesian First Aid* [77] function *bayes.t.test* by comparing pairwise differences of pools within each dataset.

### Estimation of genetic diversity and GC content

To estimate genetic diversity as π, we used Popoolation1 v.1.2.2 [74] for each pool. We used the mapped clean reads and samtools *mpileup* function to obtain mpileup files for each pool. For π estimation, we used the *Variance-sliding.pl* function with a *--min-count* of 3, a *--min-coverage* of 15, a *--max-coverage* of 50 and a *--min-covered-fraction* of 0.5. We run all analyses using a *--window-size* and *--step-size* of 1000 to avoid overlapping windows.

We estimated GC content with a custom Python script (Supplementary material) using a window size of 10kb. Because the script required the pool in .fasta format, we previously split each .bam file by chromosome and converted them to .fasta format using samtools *view* and samtools *fasta* respectively. To test whether recombination rates is correlated to π or GC content, we used again the function *bayes.cor.test* in the R package *Bayesian First Aid* [77].

### Genetic differentiation

For the experimental datasets and the time series dataset we calculated the genome-wide F_ST_ using Popoolation2 v.1.201 *fst-sliding.pl*. We looked at F_ST_ pairs between the 1^st^ and 7^th^ generation microplastics (PA1G-PA7G) treatment, and the 7^th^ generation of control and microplastics treatment (CT1G-PA7G) in the microplastics dataset. For the Cadmium dataset we looked at F_ST_ pairs between the 1^st^ and 8^th^ generation Cadmium treatment (Cd1G-Cd8G) and the 8^th^ generation control and Cadmium treatment(Ct1G-Cd8G). For the temperature selection dataset we estimated the between the time series NP0 pool (ancestral population for all experimental datasets) and each of the different temperatures treatments and replicates. In addition, we calculated the F_ST_ between NP0 pool and first and last generation for the Cadmium and microplastics dataset and NP0 pool and all treatment and replicates for the temperature dataset to be included in the structure equation model analysis (discussed below).

We investigated the relationship between genetic differentiation (F_ST_) and significant shifts in recombination rates in the experimental datasets (temperature, microplastics, and cadmium) to assess how recombination rate distributions changed between control and treatment pools over time. This analysis aimed to determine whether genomic regions experiencing substantial recombination rate shifts across generations were associated with regions of high F_ST_, which we interpret as a proxy for selection. We defined regions of high F_ST_ as regions with F_ST_ values above the 95% percentile threshold. We calculated the difference (Δ) in recombination rates between the first generation control and treatment groups, as well as between the control and the last generation. For the temperature dataset, Δ was calculated between each treatment and the NP0 pool from the seasonality dataset, representing the ancestral population. To identify genomic regions with substantial shifts in recombination rates, we extracted regions where Δ exceeded the 95% percentile. The corresponding F_ST_ estimates for these regions were then obtained to examine potential associations. Statistical relationships were assessed using the *bayes.cor.test* function from the R package *Bayesian First Aid* [77].

### Estimation of TEs

As required by popoolationTE, v.1.10.03 [78] we first prepared a custom made library a of *C.riparius* specific transposable elements and repeats with RepeatModeller v.2.0 [79] and -engine NCBI. We used the headers of each of the sequencers in the library to create a TE hierarchy, an additional requirement from popoolationTE, extracting insert name, family and order. We obtained a TE reference genome by combining the TE library and the reference genome. We mapped paired-end sequence reads of each pool from each dataset to the TE refence genome using BWA alignment algorithm *bwasw* [73]. Then we converted the resulting .sam files to bam files with popoolationTE function se2pe. We generated a mpileup file with all bam files per dataset with popoolationTE function *ppileup*, with a min quality --map-qual 15. To avoid biased comparison of TE abundance between pools within each dataset, we used the popoolationTE function *subsampleppileup* with target-coverage 55 to obtain a uniform coverage and thus, homogenize the power to identify TE insertions. In order to estimate population frequency of TE insertions and rearrangements we used the popoolationTE functions *identifySignatures* and *frequency*.

### Structural equation models

For the three experimental datasets we tested hypothesized causal relationships among different genomic parameters in order to explore the multivariate relationships among them through a series of structural equation models (SEMs). We included recombination rates, genetic diversity and GC content as delta (Δ) differences between the parameter of a particular pool with the NP0 pool from the time series dataset (ancestral pool). We used the Δ of the parameters to provide insights into how the treatment alters the relationships and dynamics between these parameters, rather than just describing their existing associations. We also include F_ST_ estimations as a pairwise estimation between a pool and the NP0 pool. Causal relationships included the impact of recombination rates and F_ST_ on genetic diversity. We used here F_ST_ as a proxy for selection, which in addition has an impact on genetic diversity. We performed the SEMs using piecewiseSEM v.2.3.0 [80] in R for all experimental datasets. We used established causal and covariance relations as shown in Figure 1. Model fit and conditional independence claims among nodes for each pool was evaluated using Fisher’s C value, with *p*-values showing statistically significance between model and data.

**Figure 1.**
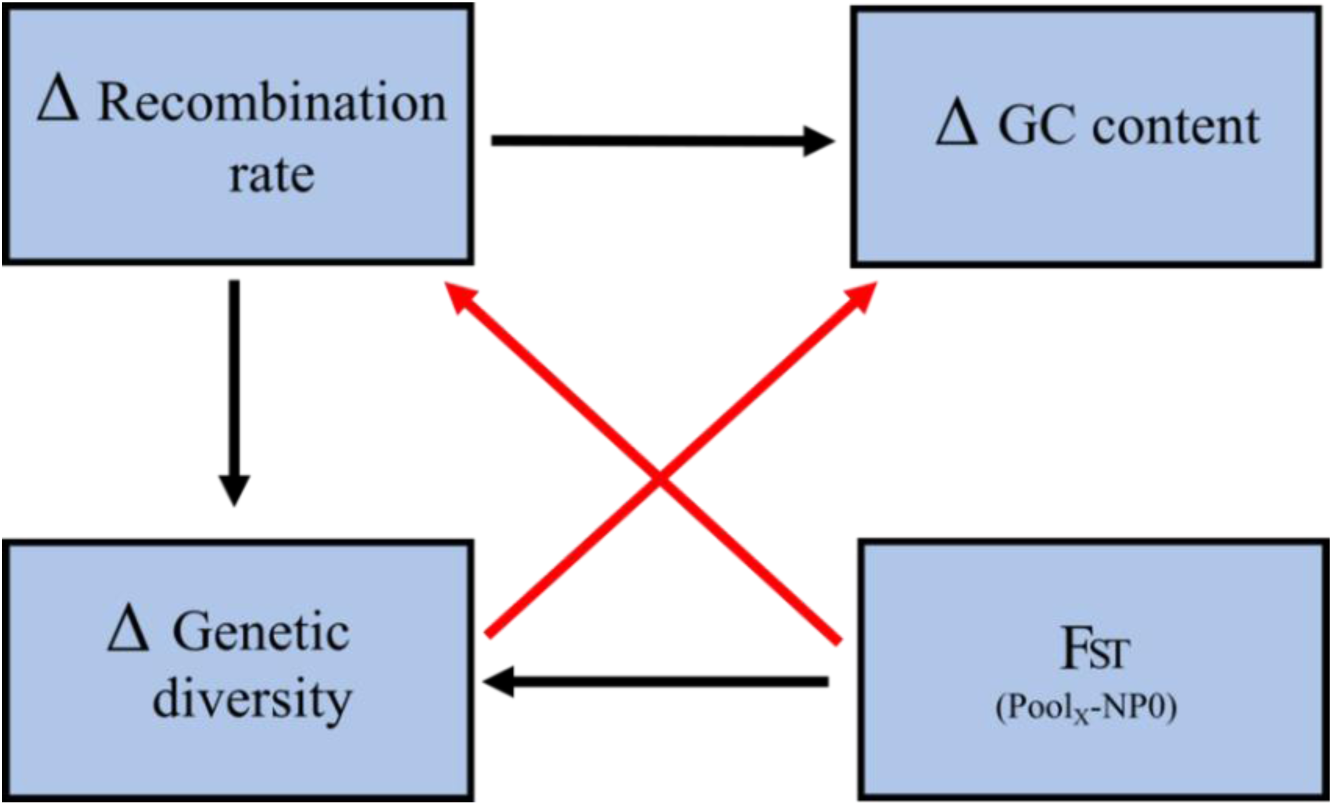
Structural equation model (SEM) of hypothesised causal and covariance relationships tested in this study for the experimental selection datasets: temperature, microplastic and Cadmium. Black arrows indicate a causal relationship and red arrows indicate a covariance relationship. Δ indicates the difference between a particular pool and NP0 pool from seasonality dataset (ancestral pool). F_ST_ corresponds to the estimation between a particular pool and NP0.

## RESULTS

### Recombination rates estimation

Mean recombination rates varied across datasets and pools within each dataset. For the geography datasets the Hessen and Rhone-Alpes pools show the highest recombination rates (2.84e-09 ± 3.60E-09 c/bp and 2.81e-09 ± 3.26E-09 c/bp) and Andalucia showed the lowest recombination rate (1.32e-09 ± 1.71e-09 c/bp). In the seasonality dataset the NP5 pool (autumn 2018) showed the highest mean recombination rate (4.78e-09 ± 3.68E-09 c/bp) followed by the cold snap pool NP5IA (winter March 2018 - 4.38e-09 ± 3.21E-09 c/bp), whereas NP4 (winter 2018) showed the lowest (2.16e-09 ± 2.59E-09 c/bp). Mean values across different pools in the selection temperature dataset showed minimal variation. Both replicates of the 20 °C treatment pools showed the lowest and highest recombination rates (ST20A=5.77e-09 ± 3.63E-09 c/bp and ST20B=6.92e-09 ± 3.98E-09 c/bp). In the selection microplastic dataset the polyamid (PA) treatment seventh generation (PA7G) pool showed the lowest recombination rate (1.85e-09 ± 2.24E-09 c/bp) and control first generation (CT1G) showed the highest (3.46e-09 ± 3.52E-09 c/bp). Pools under PA treatment showed lower recombination rates as their respective control pool. In the selection cadmium dataset the lowest recombination rates for all replicates were the Cadmium 8^th^ generation treatment. The pools with the highest recombination rate in all replicates were the Cadmium treatment pools of 1^st^ generation (Table 1).

**Table 1.** Summary of recombination rates of all pools; π estimation in 10kb windows; correlation between recombination rates and π; GC content in 10kb windows; correlation between recombination rates and GC-content; and number of TEs insertions per pool. Detailed explanation of datasets and pools IDs can be found in Figure S1.

| *Populations* | *Agent* | *Pool's ID* | *rr (c/bp)* | $\pi$ | *Corr. rr- $\pi$* | *GC content* | *Corr. rr-GC* | *# of TEs* |
| --- | --- | --- | --- | --- | --- | --- | --- | --- |
| *Natural* | *Geography* | Andalucia | 1.32E-09 | 0.0099 | 0.27 | 30% | NO | 57 |
|  |  | Piemonte | 2.06E-09 | 0.0101 | 0.25 | 29% | NO | 56 |
|  |  | Lorraine | 1.72E-09 | 0.0097 | 0.31 | 31% | NO | 56 |
|  |  | Rhone-Alpes | 2.81E-09 | 0.0111 | 0.27 | 31% | NO | 57 |
|  |  | Hessen | 2.84E-09 | 0.0099 | 0.20 | 31% | NO | 58 |
|  | *Seasonality* | NP0 | 3.07E-10 | 0.0116 | 0.40 | 32% | NO | 76 |
|  |  | NP2 | 2.44E-09 | 0.0028 | 0.35 | 31% | NO | 79 |
|  |  | NP3 | 5.37E-09 | 0.0053 | 0.33 | 30% | NO | 83 |
|  |  | NP4 | 2.16E-09 | 0.0120 | 0.20 | 31% | NO | 78 |
|  |  | NP5IA | 4.38E-09 | 3.04E-05 | NO | 31% | NO | 76 |
|  |  | NP5 | 4.78E-09 | 0.0115 | 0.29 | 32% | NO | 111 |
| *Experimental selection* | *Temperature* | ST14A | 6.30E-09 | 0.0112 | NO | 31% | NO | 93 |
|  |  | ST14B | 6.33E-09 | 0.0111 | moderate (0.15) | 31% | NO | 57 |
|  |  | ST20A | 5.77E-09 | 0.0111 | moderate (0.19) | 31% | NO | 57 |
|  |  | ST20B | 6.92E-09 | 0.0105 | 0.21 | 31% | NO | 64 |
|  |  | ST26A | 6.46E-09 | 0.0093 | moderate (0.10) | 32% | NO | 56 |
|  |  | ST26B | 6.48E-09 | 0.0004 | 0.34 | 30% | NO | 61 |
|  | *Microplastics* | CT1G | 3.46E-09 | 0.0048 | 0.40 | 28% | NO | 218 |
|  |  | PA1G | 2.51E-09 | 0.0061 | 0.21 | 28% | NO | 224 |
|  |  | CT7G | 3.12E-09 | 0.0064 | 0.58 | 29% | low moderate (0.11) | 363 |
|  |  | PA7G | 1.85E-09 | 0.0083 | 0.42 | 29% | NO | 194 |
|  | *Cadmium* | CtA1G | 1.22E-09 | 0.0004 | 0.25 | 30% | NO | 305 |
|  |  | CtA8G | 1.72E-09 | 0.0020 | NO | 30% | NO | 251 |
|  |  | CtB1G | 2.08E-09 | 0.0003 | 0.25 | 20% | NO | 265 |
|  |  | CtB8G | 1.93E-09 | 0.0012 | 0.37 | 30% | NO | 263 |
|  |  | CtC1G | 8.84E-10 | 0.0004 | 0.22 | 29% | low moderate (0.1) | 230 |
|  |  | CtC8G | 8.45E-10 | 0.0014 | 0.36 | 29% | NO | 227 |
|  |  | CdA1G | 2.95E-09 | 0.0004 | 0.28 | 30% | NO | 231 |
|  |  | CdA8G | 8.95E-10 | 0.0012 | 0.26 | 30% | NO | 209 |
|  |  | CdB1G | 2.19E-09 | 0.0004 | 0.24 | 30% | low moderate (0.11) | 239 |
|  |  | CdB8G | 8.96E-10 | 0.0006 | NO | 29% | NO | 227 |
|  |  | CdC1G | 1.95E-09 | 0.0004 | 0.26 | 20% | NO | 205 |
|  |  | CdC8G | 1.74E-09 | 0.0019 | 0.38 | 30% | NO | 232 |

For the geography, selection temperature and microplastics datasets, chromosome 4 in all pools showed the highest recombination rate. In the seasonality dataset, pools NP0 and NP2 showed the highest recombination rates in chromosome 4 and, pools NP3, NP4, NP5IA and NP5 in chromosome 3. Eight out of the 12 pools from the selection Cadmium dataset showed the highest recombination rates in chromosome 4, while the remaining pools showed the highest recombination rates in either chromosome 1, 2 or 3.

Distribution of 10kb averages recombination rates along the genome showed the decrease of the rates at the start and end of chromosome (Figure S2a-e). The genome-wide distribution of recombination rates in the Hessen pool, compared with experimental datasets, revealed significantly lower recombination rates in the Hessen pool than in the temperature dataset pools. This pattern was not as clearly observed in the microplastics and cadmium dataset pools (Figure S3).

Within each dataset pool pairwise comparisons showed that all observed differences amongst pools were different from zero with the exception of replicates A and B for the 26 °C treatment of the temperature dataset and control first generation and Cadmium first generation of replicate B of the Cadmium dataset (Table 1).

Comparison of Hessen pool from the climate dataset, from which the ancestral source population for the selection temperature, microplastics and Cadmium datasets pools come from, are shown in Figure 2 and Figure S3.

**Figure 2.**
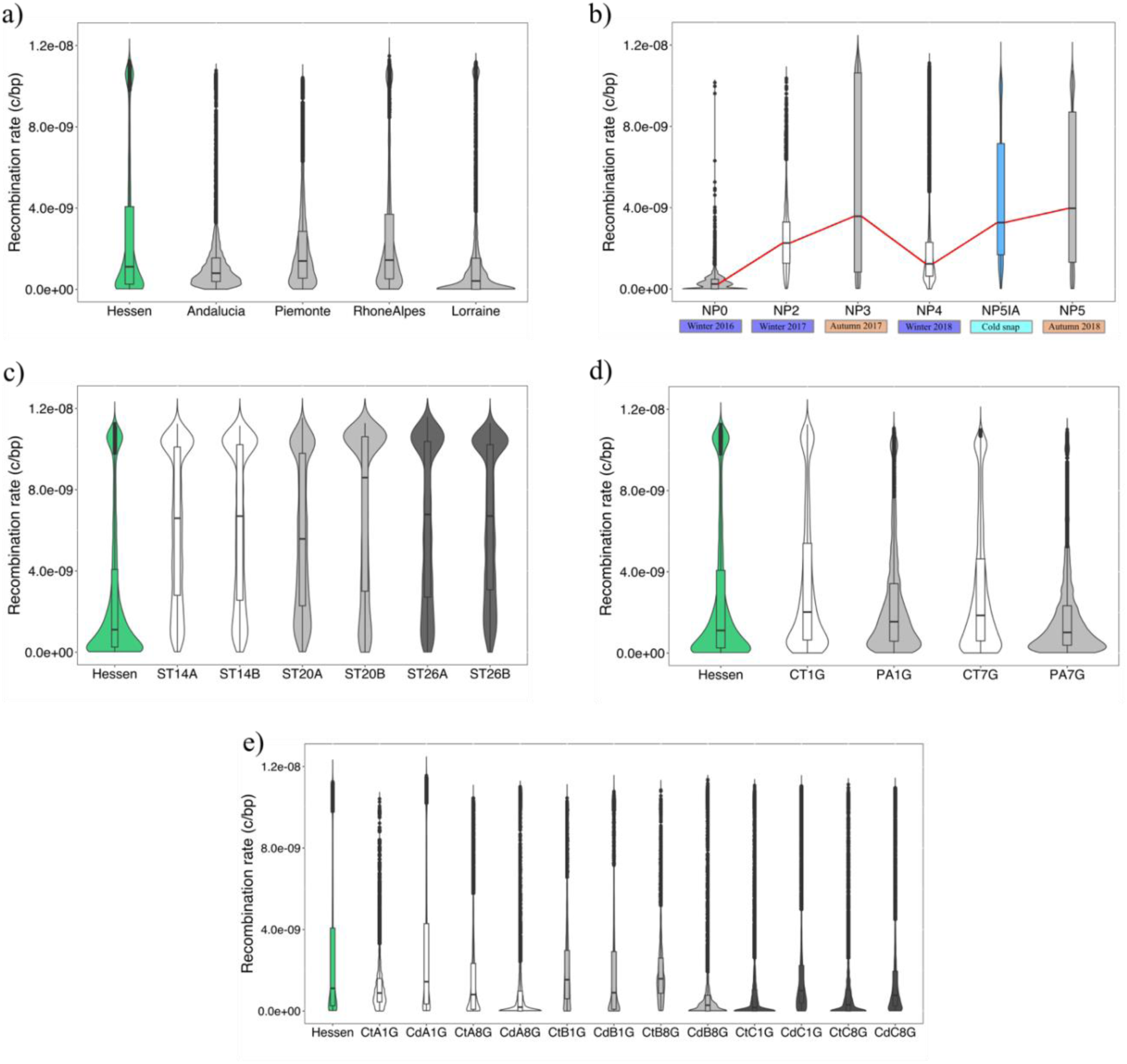
Recombination rates comparison of natural populations datasets: (a) geography and (b) seasonality; and experimental selection datasets: (c) temperature, (d) microplastics and (e) Cadmium datasets. For the experimental selection datasets populations the ancestral population (Hessen) is shown for comparison.

### Differences in genetic diversity, GC content and TEs across pools

π mean average estimations were generally low for all pools and range from 3.04e-05 in the for the cold snap pool (NP5IA) to 0.012045 from the 2018 winter pool (NP4) from the seasonality dataset (Figure 3 and Table 1). Pairwise differences were statistically significant, however a relevant effect size was only observed between 1^st^ and 7^th^ generation of the PA treatment of the microplastic dataset, between the different temperature treatments in replicate B of the temperature selection dataset and between the control and Cd treatment of 8^th^ generation for all replicates in the Cd dataset.

**Figure 3.**
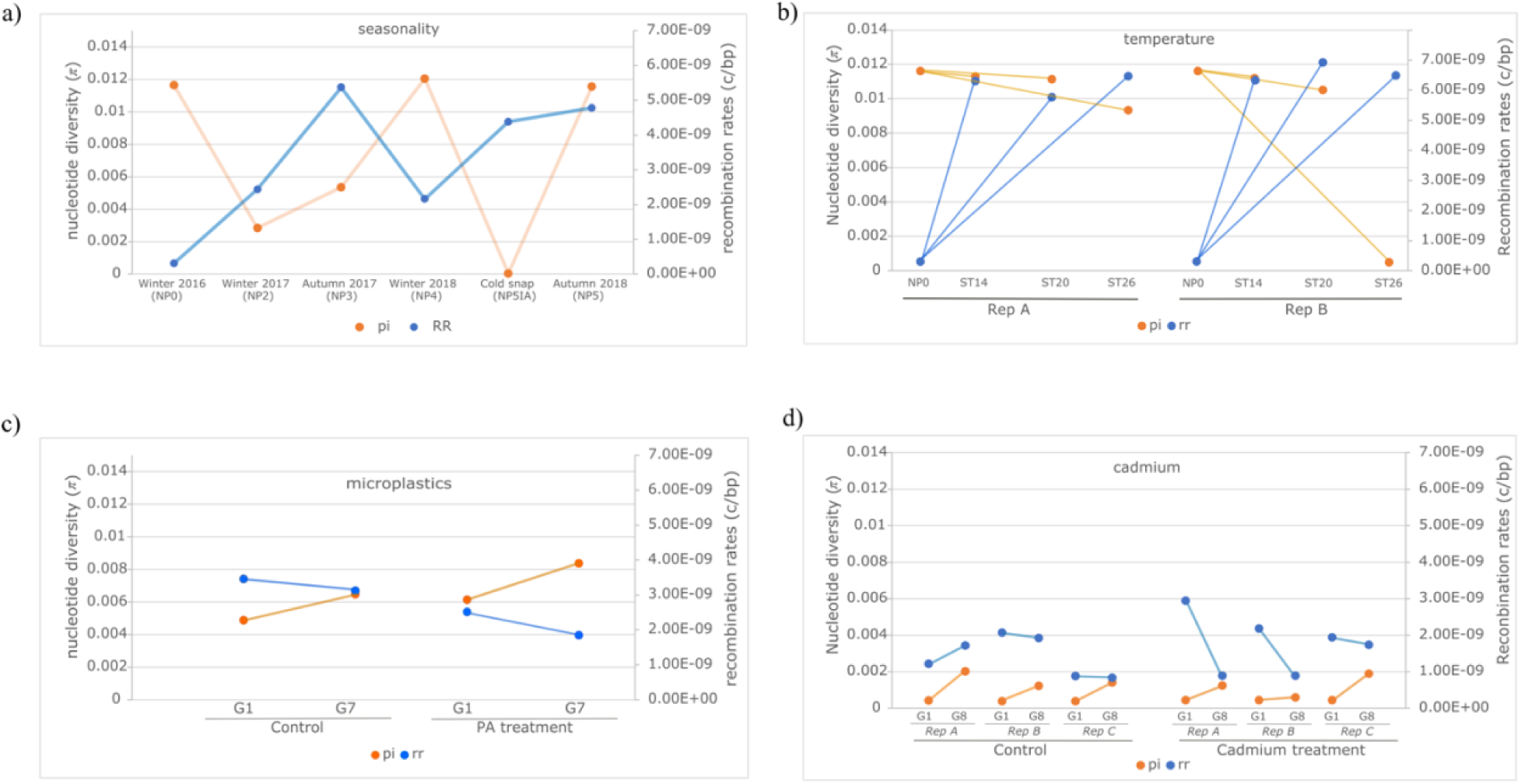
Mean genetic diversity (π; primary Y axis) and mean recombination rates estimates (secondary Y axis) of a) seasonality dataset; b) temperature dataset. Rep= replicate. NP0=ancestral pool from seasonality dataset (NP0), ST14, ST20 and ST26 refers to the different temperature treatments, 14°C, 20°C and 26°C, respectively; c) microplastics dataset. G1=generation 1 and G7=generation 7; and d-f) Cadmium dataset. Rep= replicate. G1=generation 1 and G8=generation 8.

GC content average estimation range from 0.28899 for the Control 1^st^ generation of the microplastics dataset to 0.322158 for the Autumn 2018 pool (NP5) from the seasonality dataset (Table 1). Pairwise differences were statistically significant, however significant effect size was only observed between 1^st^ and 7^th^ generation of the control and PA treatment of the microplastic dataset, between the 14°C and 26°C, and 20°C, and 26°C, of both replicates in the temperature selection dataset, between the control and Cd treatment of 1^st^ generation for replicate C in the Cd dataset, and between all pools in the time series dataset.

The abundance of TEs in pools, as measured by the count of TEs in the genomes, range from 56 in several pools (from geography and selection temperature datasets) to 363 (in microplastics dataset). Furthermore, average number of TEs of the microplastics and Cadmium datasets was 3.5 times higher than average number of TEs of the climate, seasonality and temperature selection dataset (Table 1).

### Statistical analyses

#### Correlation between recombination rates and different genomic parameters

Bayesian pair test showed generally a strong positive association between recombination rates and π for all pools except for the cold snap pool from the seasonality dataset (NP5IA), replicate A of the 14 °C treatment in the temperature dataset, and control replicate A 8^th^ generation and Cd treatment replicate B 8^th^ generation from the Cadmium dataset (Table 1).

Bayesian pair tests also showed no positive association between recombination rates and GC content for all pools except for low moderate correlation for pools control 7^th^ generation in microplastics dataset and, control replicate C 1^st^ generation and Cd treatment replicate B 1^st^ generation in the Cadmium dataset (Table 1).

We also identified genomic regions for which the Δ in recombination rates showed a skewed distributions in replicate A and B of the Cadmium dataset and, replicate A and B of the temperature dataset (Figure 4). In particular, the distribution the Δ in recombination rates for the A and B of the Cadmium treatment extends more into negative values compared to the control Δ distribution (Figure 4 a-b), implying that in some genomic regions, recombination rates decreased more under Cadmium treatment than in the control. Replicate C of the Cadmium dataset and pools from the microplastics datasets did not present this pattern (Figure S4).

**Figure 4.**
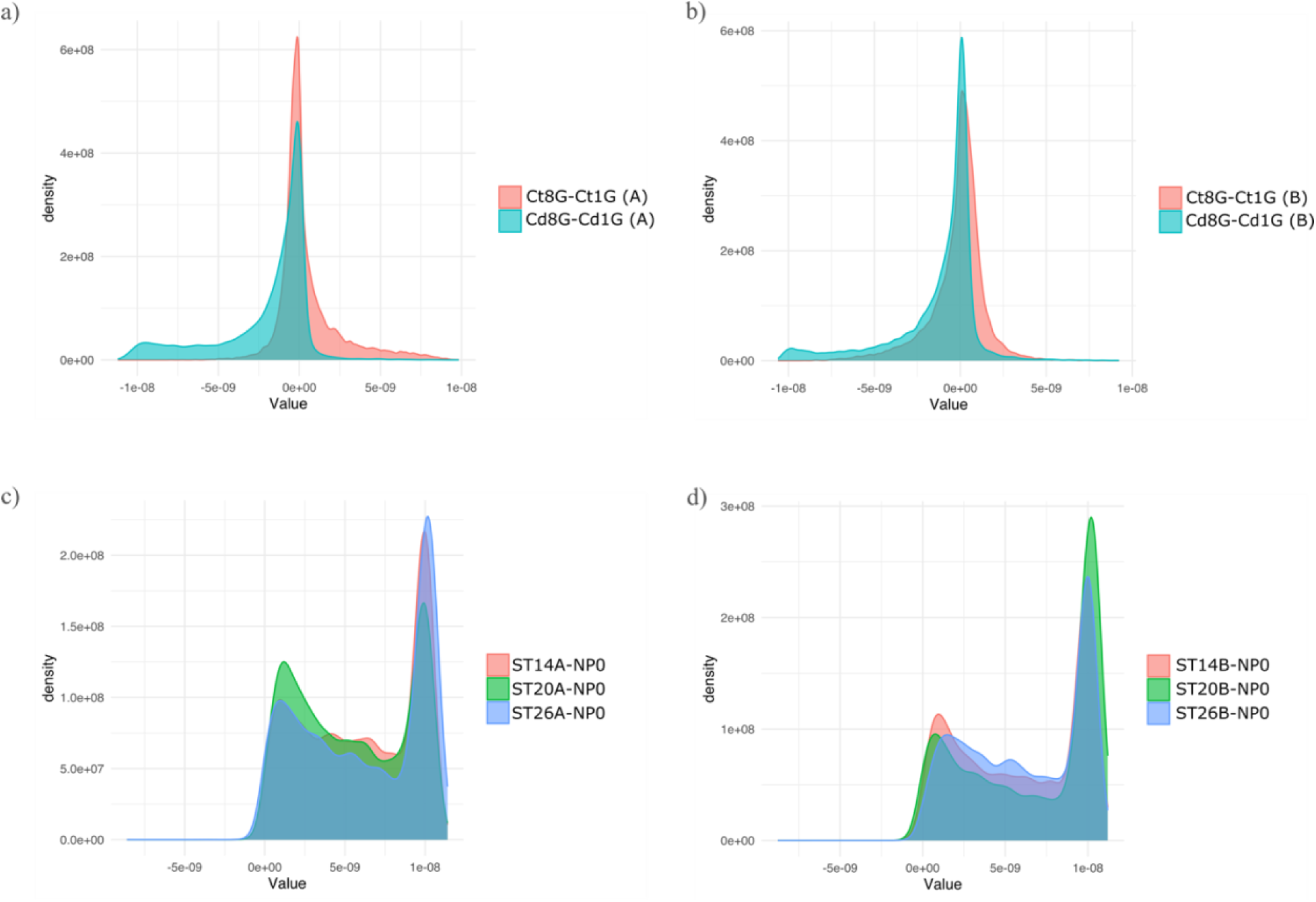
Plots showing the distribution of difference (Δ) in recombination rates between: a) 8^th^ and 1^st^ generation for control (pale red) and Cadmium (cyan) pools replicate A; b) 8^th^ and 1^st^ generation for control (pale red) and Cadmium (cyan) pools replicate B; c) temperature treatments and ancestral pool from seasonality dataset (NP0) in replicate A, temperature dataset; and d) temperature treatments and ancestral NP0 in replicate B.

We found positive correlation between regions with high Δ in recombination rates and their corresponding F_ST_ estimates for those regions for A and B of the Cadmium treatment (ρ = 0.10 and 0.082, respectively; Figure S5a). 95% HDI for both correlations do not include zero. However, we found no correlation for all temperature treatments in both replicates A and B, and all 95% HDI did indeed include zero (Figure S5b).

### Structural equation models

In general, the SEM model showed to be a good fit for the majority of the pools analysed. In the temperature dataset Fisher’s C showed that the model explained the data well and no other variables connections are missing, except for the replicate A 20°C treatment pool, where a relationship between GC content and F_ST_ is proposed with moderate significance (\*\*\**p* < 0.001). Causal relationship between recombination rates and genetic diversity and F_ST_ with genetic diversity are statistically significant for all pools except for replicate A 14°C treatment and replicate B 26°C treatment, respectively. Causal relationship between recombination rates and GC content is only statistically significant for replicate A 26°C treatment and replicate B 14°C treatment (Table S2). We also detected a reduction in the estimates of the covariance between F_ST_ and recombination rate as temperature increased from 14°C to 26°C. (Figure S6a). In the microplastics dataset Fisher’s C showed that the model explained well the data and no other variables connections were missing. All hypothesized causal relationships are statistically supported with high support (\*\*\**p* = 0; Figure S6b). Covariance relationships are only supported for control 7G pool between F_ST_ and recombination rates with moderate support (\**p* < 0.01). In the replicate A of the Cadmium dataset all hypothesized causal and covariance relationships were statistically supported with high support (\*\*\**p* = 0) for both pools except for the covariance between F_ST_ and recombination rates in the PA treatment 8^th^ generation (Figure S6c). In addition, an additional missing relationship is proposed between GC content and F_ST_ with high statical support (\*\*\**p* = 0; Figure S6c). In the replicate B control and PA treatment for both 8^th^ generations of the Cadmium dataset causal relationships are highly supported except for the relationship between recombination rates and GC content (Figure S6c). In addition, covariance relationships are supported except for the π and GC content in PA treatment 8^th^ generation (Figure S6c). In replicate C, results were only available for control 8^th^ generation, where causal and covariance relationships were highly supported except for causal relationship between recombination rates and GC content, and the covariance relationship between π and GC content (Figure S6c). In addition, an additional missing relationship is proposed between GC content and F_ST_ with low statical support (\**p* < 0.01; Figure S6c).

## DISCUSSION

Understanding recombination rates is crucial in evolutionary biology, as this process not only drives genetic diversity but also shapes the patterns of natural selection and adaptation across populations. In this study we showed how recombination rates for *C. riparius* vary across a latitudinal range, seasons and under different experimental treatments. As anticipated, our findings reveal substantial variability in recombination rates across the datasets analyzed.

### Variation of recombination rates in natural populations

The five populations represented in the geographic dataset span a broad range of environmental and climatological conditions, reflecting the heterogeneity of regions sampled, from southern to central Europe. Previous works on natural populations have linked diversity in climatic factors and latitudinal cline to variation in recombination rates [18, 23]. The selective pressures derived from climate variation changes continuously, as can be expected for the evolutionary response of populations along these gradients. Therefore, and taking the heterogeneity of regions sampled, it is not surprising that a clear pattern cannot be observed from our data on the relationship of global recombination rates and latitude (Figure 2a). [69] demonstrated that local climate conditions exert strong selection pressures on the same populations we studied, driving climate adaptation. A study on two ecological (xeric *vs* sub-alpine) and geographical separated populations of *D. pseudoobscura* discuss the role of natural selection explaining the differences in recombination rates (as accounted by cross-over rate) of the populations [18]. One explanation they offered is that higher temperatures in one of the populations could have caused the increased recombination rates, which is in line with the fact that recombination rates is known to be plastic with regard of ambient temperature [81]. Pettrich & Waldvogel (unpublished observations) recently used whole-genome sequencing (WGS) data from the same five geographically distant populations analyzed in this study and reconstructed genome-wide recombination landscapes, highlighting the potential influence of a transposable element – the *Cla*-element – on recombination rate variation. They found reduced recombination in annotated centromere regions and could find the tendency of the element to occur in association with lower recombination rates. In their analysis, Lorraine, Rhone-Alpes, and Piemonte populations exhibited higher intrapopulation variability given in ρ. Our findings indicate greater recombination rate variation in the Rhone-Alpes, Piemonte, and Hessen populations (Figure 2a). Furthermore, the variability in Rhone-Alpes and Piemonte align with the other study, while we find different variability for Lorraine. While both studies focused on the same natural populations, we here used different metrics, as ρ is a scaled measured of r (ρ = 4N_e_*r), and as such can be directly influenced by effective population size (*N_e_*). The ancient N_e_ for Lorraine is estimated to be lower than the other populations [72] which could lead to more variability in ρ, also considering effects caused by differences between WGS and PoolSeq data [82].

Although much research has focused on how recombination varies across spatial environmental gradients, temporal variation—particularly across seasons—remains relatively underexplored. Our data reveal a subtle but consistent seasonal fluctuation in genome-wide recombination rates in *C. riparius* (Figure 2b). Specifically, autumn pool samples (NP3 and NP5) show generally higher recombination rates compared to those collected in winter (NP0, NP2, and NP4). Several interrelated factors may contribute to this pattern. One explanation is that warmer autumn temperatures promote higher recombination rates, while colder winter conditions might physiologically suppress recombination. This pattern aligns with observations in *Drosophila melanogaster*, where recombination is both genetically and plastically reduced under lower temperatures [30]. Additionally, [33] found seasonal variation in crossover rates and interference in *D. melanogaster*, suggesting recombination can be shaped by indirect selection driven by seasonal environmental stresses. However, beyond immediate environmental effects, differences in generational turnover between seasons may also play a critical role. In *C. riparius*, generation times are temperature-dependent, with rapid turnover during warmer months (spring and summer) and a developmental pause or zero turnover during winter. Thus, autumn populations are the result of several overlapping generations accumulating over the productive summer, potentially amplifying recombination signatures through repeated meiosis. In contrast, winter samples reflect only a single or even arrested generation, leading to relatively reduced signals of recombination. This generation-count bias offers a compelling explanation that is independent of environmental plasticity. Essentially, more generations before autumn sampling means more opportunities for recombination events to shape the population’s genetic landscape. This may be further compounded by reduced *N_e_* in winter due to ecological bottlenecks, which would constrain the recombination signal both through demographic contraction and selection favoring well-adapted genotypes. Taken together, the observed seasonality in recombination rates likely reflects a complex interplay of ecological, developmental, and evolutionary processes, where both environmental plasticity and life-history dynamics influence genome-wide recombination patterns.

Furthermore, we observed that the cold snap of March 2018 (NP5IA) led to a noticeable increase in recombination rates within a short period of time compared to winter 2018 (NP4) (Figure 2b). This appears as a stress-induced response to the sudden drop in temperatures. This plastic response may help populations rapidly generate new allele combinations in response to selection pressures. Indeed, [83] demonstrated that this cold snap resulted in selection for specific polygenic traits to enhance adaptation to sudden environmental changes. Consistent with this, we detected a drastic reduction in genetic diversity in NP5IA, supporting the role of selection in shaping the observed patterns (Table 1) Since the cold snap likely occurred within a single generation, with no new recombination events taking place, selection may have directly influenced recombination rate estimates. Alternatively, it is possible that only individuals with higher recombination rates in previous generations survived, as suggested by our results. (Figure 2b).

### Experimental selection impact on recombination rates

Our results reveal a complex pattern of recombination rate variation across experimental treatments in the temperature dataset, with distinct but opposing trends observed between the two replicates. However, the different thermal regimes do not seem to have a high impact on recombination rates (Figure 2). In replicate A, recombination rates followed a subtle U-shaped pattern across increasing temperature treatments, a pattern observed in *D. melanongaster* [84], while in replicate B, an inverted U-shaped pattern emerged. Notably, we established that these patterns are not driven by temperature regimes themselves but rather by adaptation to laboratory conditions. Evidence from [85], points to selection process due laboratory conditions, in particular selection on traits related to conductivity tolerance and developmental speed, rather than to the different temperature regimes. Despite these contrasting recombination trajectories, both replicates exhibited a consistent decline in genetic diversity.

The opposing recombination trends between replicates suggest that selection under laboratory conditions does not impose a uniform effect on recombination across populations but instead interacts with pre-existing genetic variation and stochastic evolutionary processes. One potential explanation is that recombination rate variation is influenced by genetic background differences between the replicates. If different initial standing genetic variation affects recombination modifiers, selection could drive distinct trajectories in each population. Additionally, recombination itself may evolve as a response to laboratory adaptation, with selection favoring different recombination dynamics depending on the genomic architecture of each replicate.

The concurrent decrease in genetic diversity across all samples suggests that selection has played a dominant role in shaping the genomic landscape, leading to the loss of allelic variation. This supports the hypothesis that adaptation to laboratory conditions has imposed strong selection pressures, reducing neutral diversity while allowing beneficial alleles to rise in frequency. However, the decoupling of recombination rate patterns and genetic diversity further indicates that recombination does not uniformly facilitate or constrain adaptation but instead interacts with selection in a more nuanced manner.

Our analysis also revealed distinct genomic regions exhibiting exceptionally high recombination rates across all six experimental populations, regardless of temperature treatment. Given that prior evidence [85] indicates strong selection for laboratory adaptation rather than direct temperature effects, these elevated recombination rates are likely linked to the selective pressures imposed by the lab environment. If increased recombination confers an adaptive advantage in the novel laboratory environment—by accelerating beneficial allele combinations—then selection may favor alleles that promote higher recombination rates in specific genomic regions. This mechanism would result in persistent regions with high recombination even after adaptation has progressed over multiple generations (22, 44 and 65 generations for the 14°C, 20°C and 26°C treatments, respectively).

Interestingly, we found no correlation between genomic regions with high Δ recombination rates (i.e., recombination rate differences between each pool and the ancestral pool) and F_ST_ estimates for the same genomic regions (Figure 2 and Figure S5b). This suggests that recombination rate changes are not directly associated with strong allele frequency differentiation between treatment and ancestral populations. However, interpreting this lack of correlation requires caution due to potential technical and biological caveats. Specifically, regions of high F_ST_ often coincide with areas of reduced nucleotide diversity (π), consistent with selective sweeps. Because most recombination rate inference tools—including those used in this study—rely on the presence of informative haplotypes or segregating variation, recombination is inherently more difficult to detect in regions where diversity has been eroded by strong selection. As such, apparent “low recombination” in high-F_ST_ regions may partly reflect reduced detectability, rather than an absence of recombination per se. This issue is well documented, for example, in vertebrate breeding populations, where low genetic diversity hampers the tracking of recombination events across generations. Thus, while our findings still suggest that recombination rate shifts may not be tightly coupled with F_ST_-based differentiation, it is also possible that part of the recombination signal is masked in sweep regions. This underscores that the observed recombination patterns may represent only a subset of the true genomic landscape, with high-confidence estimates biased toward regions of moderate diversity. It is worth emphasizing that selection may still act to increase recombination in specific loci—either to facilitate adaptation by breaking linkage with deleterious alleles, or as a trait under selection itself—without producing strong F_ST_ signals, especially if the underlying adaptive variants derive from standing genetic variation. Overall, the absence of a strong correlation between recombination shifts and F_ST_ should be interpreted as a nuanced signal: one that potentially reflects both biological mechanisms and analytical limitations tied to diversity-dependent recombination inference.

Regarding changes of recombination rates in relation to non-temperature stressors [86, 87], our study shows that exposure to both microplastics and Cadmium, impacts recombination rates. As observed for the temperature selection dataset, there is a significant increase in recombination rates in the microplastics datasets in comparison to the ancestral population from Hessen (Figure 2d). Surprisingly, we found that exposure to microplastics as a potential environmental stressor did not lead to an increase in recombination rates, neither in the first nor the seventh generation. Instead, we observed a genome-wide reduction in recombination rates in the microplastics treatment pools compared to the control pools, with this reduction becoming even more pronounced by the seventh generation (Figure 2d). This pattern may be explained by three possible mechanisms. One possible explanation is that stress-induced DNA damage may trigger rapid repair mechanisms that prioritize genome integrity over generating genetic diversity through recombination. In such cases, cells may favor non-homologous end joining (NHEJ), a faster but more error-prone repair pathway, over homologous recombination, which is typically associated with meiotic recombination and genome reshuffling [88–91]. Indeed, [92] and [93] that microplastic exposure induces oxidative stress and identified genes under selection associated with oxidative stress pathways. Similarly, [69] found that candidate loci for local adaptation in their study were enriched for genes involved in detoxification processes, such as transport, phosphorylation, and immune responses. These gene ontology (GO) terms, commonly linked to stress responses, have also been identified in *D. melanogaster*, suggesting conserved adaptive mechanisms across species in response to environmental pressures [94–96].

Secondly, a reduction in recombination may help preserve beneficial genetic combinations already present in the population. By limiting recombination, the genome may prevent the breakup of advantageous allelic associations, which could be particularly beneficial in stable or stressful environments where maintaining successful genotypes provides a selective advantage [97–99]. This genetic conservatism may be especially relevant in laboratory-maintained populations, where pre-adaptation to controlled conditions could have occurred due to long-term isolation from the original wild populations. Finally, it is possible that one of the selected loci is linked to or epistatically correlated with a recombination-modifying locus, such that selection for a trait that mitigates the effects of microplastic exposure inadvertently reduces recombination rates [19].

Interestingly, we observed a temporal increase in π in both control and microplastic-treated pools, despite the decrease in recombination (Figure 3). This suggests that the observed increase in genetic variation is not due to recombination but rather to elevated mutation rates. Previous studies have shown that environmental stressors can drive extreme mutation rates, introducing novel genetic variation into populations even within five generation in *C. riparius* [100, 101]. In addition, exposure to metals such as copper and nickel has been linked to increased mutation rates in *Daphnia pulex*, primarily through elevated rates of gene deletions and duplications [102]. A similar mechanism may be at play in our study, where microplastic-induced oxidative stress induced the production of reactive oxygen species (ROS), which in turn could be accelerating mutation accumulation. Oxidative DNA lesions, such as 8-oxoguanine, are known to introduce replication errors, leading to an increase in point mutations and small insertions or deletions (indels) [103]. If these mutations are not efficiently repaired or are selectively tolerated due to stress-adaptive pressures, they could result in a net increase in genetic diversity. Together, these findings suggest that microplastic exposure leads to reduced recombination but elevated mutation rates, likely as a consequence of oxidative stress-induced DNA damage, which has significant implications for the long-term evolutionary trajectories of populations exposed to environmental stressors.

Our findings indicate that recombination rates were generally higher in control samples compared to Cadmium-treated populations, except for replicate A in the first generation and both generations in replicate C. This suggests that exposure to Cadmium may suppress recombination over multiple generations, possibly due to stress-induced DNA damage leading to a shift toward NHEJ over homologous recombination as discussed above for microplastics. However, the exceptions observed in replicate A in the 1^st^ generation and replicate C, for both generations, suggest that additional factors, such as genetic background, stochastic effects, or differences in selective pressures, may be influencing recombination responses on a population-level. The overall reduction in recombination rates under Cadmium exposure aligns with previous studies showing that toxic substances can lead to a more conservative genome structure, potentially limiting genetic reshuffling and affecting long-term adaptive potential ([35] - DDT; [104] - atrazine; [105–108] - Cadmium).

Patterns of genetic diversity also varied across replicates and generations, reflecting distinct evolutionary responses to Cadmium exposure. While genetic diversity remained similar between control and Cadmium-treated samples in the first generation, divergence emerged by the eighth generation. In replicates A and B, control populations exhibited higher genetic diversity, whereas in replicate C, genetic diversity was greater under Cadmium treatment. This variation suggests that while selection and genetic drift influence population differentiation, the balance between mutation, recombination, and selection differs among replicates. Furthermore, the observed positive correlation between large delta differences in recombination rate (Cadmium 8th generation vs. Cadmium 1st generation) and F_ST_ from an ancestral population in replicates A and B—but not in replicate C—suggests that recombination suppression may be linked to increased genetic differentiation from the ancestral population. This could indicate that in some populations, reduced recombination constrains genetic exchange, reinforcing divergence due to selection or drift, while in others, alternative mechanisms such as mutation-driven adaptation play a larger role.

The notably elevated number of TEs observed in both control and exposed samples within the Cadmium and microplastics datasets may be attributed to a combination of prolonged laboratory maintenance and the associated effects of reduced genetic diversity, rather than direct exposure to toxicants alone. While genotoxic stress from cadmium and microplastics is known to induce DNA damage and oxidative stress — factors that can compromise TE silencing pathways (e.g., DNA methylation, piRNA; [60, 109]) — the lack of a clear difference between treated and control groups suggests an alternative explanation. It is likely that these populations were kept in benign laboratory conditions for more generations than other datasets, providing more time for TE proliferation without selective constraints. In small, bottlenecked populations typical of lab rearing, purifying selection is often relaxed, and genetic drift may reduce the effectiveness of TE suppression mechanisms. Over multiple generations, this can lead to the accumulation of TE insertions, even in the absence of continued environmental stress. Additionally, TE expansion is a gradual process that may not manifest rapidly following acute exposure but instead becomes pronounced over time. These findings support the idea that both environmental history and experimental design—especially time in the lab—are critical factors influencing TE dynamics.

### Recombination dynamics and its causal relationships

The SEMs applied across the three experimental datasets provided valuable insights into the causal relationships among recombination rates, genetic diversity, GC content, and F_ST_, highlighting both shared patterns and dataset-specific variations.

In the temperature dataset, our findings indicate as expected that recombination rates and F_ST_ exert significant causal effects on genetic diversity in most pools, suggesting that changes in recombination frequency and selection pressures shape genome-wide genetic variability. However, the absence of statistically significant relationships in specific treatments, such as the 14°C in replicate A and the 26°C in replicate B, implies that the effects of temperature on these parameters may be context-dependent, potentially influenced by additional factors such as selection pressures derived from the lab conditions (Figure S6a). Interestingly, we observed a reduction in covariance estimates between F_ST_ and recombination rates as temperature increased, suggesting a potential link between changes in F_ST_ and alterations in recombination rates. However, as previously noted, temperature regimes do not directly influence the overall magnitude of recombination rates. Instead, the observed decrease in covariance appears to be associated with the number of generations exposed to higher temperatures. Specifically, a stronger relationship is evident at 14°C (22 generations), which gradually wears off by 26°C (65 generations).

In the Cadmium dataset, results varied across replicates, reflecting complex interactions between heavy metal stress and genomic responses. Replicate A showed strong statistical support for all hypothesized causal and covariance relationships, indicating that recombination rates, F_ST_, and genetic diversity were tightly interconnected under Cadmium exposure. The emergence of an additional causal pathway between GC content and F_ST_ suggests that Cadmium may exert selective pressures that influence nucleotide composition, possibly due to biased mutation processes or differential repair efficiency. In replicate B, the general pattern of causality was maintained, though the recombination-GC content relationship was not supported, suggesting that while recombination and GC content are often linked, their relationship may be disrupted under Cadmium stress, potentially due to metal-induced alterations in DNA methylation or repair mechanisms. Interestingly, in replicate C, results were only available for the control 8th generation, where most hypothesized relationships were supported except for the recombination-GC content causal link and the covariance between genetic diversity and GC content. The presence of a low-support additional relationship between GC content and F_ST_ further highlights potential stress-induced shifts in nucleotide composition. Overall, the differences in SEM outcomes across Cadmium replicates suggest that while recombination rates and selection (F_ST_) consistently shape genetic diversity, the specific interactions between genomic parameters may vary depending on lineage-specific adaptive responses or experimental variation. The observed differences in causal pathways across replicates underscore the complexity of evolutionary responses to heavy metal exposure and highlight the necessity of multi-replicate designs to capture the full spectrum of potential genomic adaptations to environmental stressors.

The microplastics dataset exhibited a more consistent pattern, with all hypothesized causal relationships being strongly supported. The statistical significance of all modeled pathways reinforces the idea that microplastic-induced stress may exert a stable and predictable influence on recombination rates, genetic diversity, and GC content. The weaker support for covariance between F_ST_ and recombination rates, observed only in the control 7G pool, suggests that the impact of microplastics on these genomic parameters is driven more by direct effects than by co-variation (Figure S6c). Given that microplastics have been associated with oxidative stress and DNA damage, it is possible that the consistent causal relationships observed in our SEM models stem from stress-induced genomic instability, which in turn influences recombination dynamics and subsequent patterns of genetic diversity. These results suggest that environmental contaminants like microplastics may exert stronger and more uniform selective pressures compared to temperature, potentially leading to predictable genomic responses across different experimental replicates. It is important to note that the microplastics dataset is based on significantly fewer generations compared to the temperature dataset. Therefore, it is possible that selection initially acts on other traits before influencing recombination rates.

## CONCLUSIONS

Recombination rates varied substantially among natural populations, suggesting that local adaptation and environmental heterogeneity play a role in shaping recombination landscapes. While a clear latitudinal pattern was not evident, differences in recombination rates among sampling locations may reflect adaptation to regional conditions. We also detected subtle seasonal variation in recombination rates, with autumn populations exhibiting higher rates than winter populations, indicating that recombination can respond to temporal environmental fluctuations. Furthermore, our experimental evolution data revealed that both microplastic and cadmium exposure can influence recombination rates, potentially as a consequence of selection favoring specific genetic combinations. These findings highlight the dynamic nature of recombination and its responsiveness to a range of evolutionary and environmental factors over generations. Our study contributes to a growing body of evidence demonstrating that recombination rate variation is a complex temporally variable in the short-term phenomenon influenced by multiple interacting factors, including genomic features, environmental conditions, and selective pressures. Further research is needed to fully elucidate the specific mechanisms driving recombination rate variation in natural populations and to explore the long-term evolutionary consequences of altered recombination rates in changing environments.

## LIST OF ABBREVIATIONS

AF: allele frequency
c/bp: centimorgan/base pair
CI: confidence interval
DDT: dichlorodiphenyltrichloroethane
ENA: European nucleotide archive
IDs: identifiers
*F*_ST_: fixation index
GO: gene ontology terms
HDI: highest density interval
NCBI: National Center for Biotechnology Information
*N_e_*: effective population size
NHEJ: non-homologous end joining
PA: polyamide
ROS: reactive oxygen species
rr: recombination rate
SEMs: structural equation models
TE: transposable elements
WGS: whole-genome sequencing

## DECLARATIONS

### Ethics approval and consent to participate

Not applicable

### Consent for publication

Not applicable

### Availability of data and materials

Raw data used in this study can be found under ENA project numbers: PRJEB19848, PRJEB35534, PRJEB32795, PRJEB48137 and PRJEB90147.

### Author’s contributions

MENB and MP designed the study. CC performed mapping. MENB collected data and performed analyses. LP assisted with TEs analyses interpretation. MENB wrote the manuscript and all authors contributed to the preparation of the final version and approved it.

## Acknowledgements

We thank M. Balint for advice on SEMs analyses. We thank anonymous colleagues for their feedback on an earlier drafts of this manuscript.

## Supplementary Material and Methods

**Table S1.**
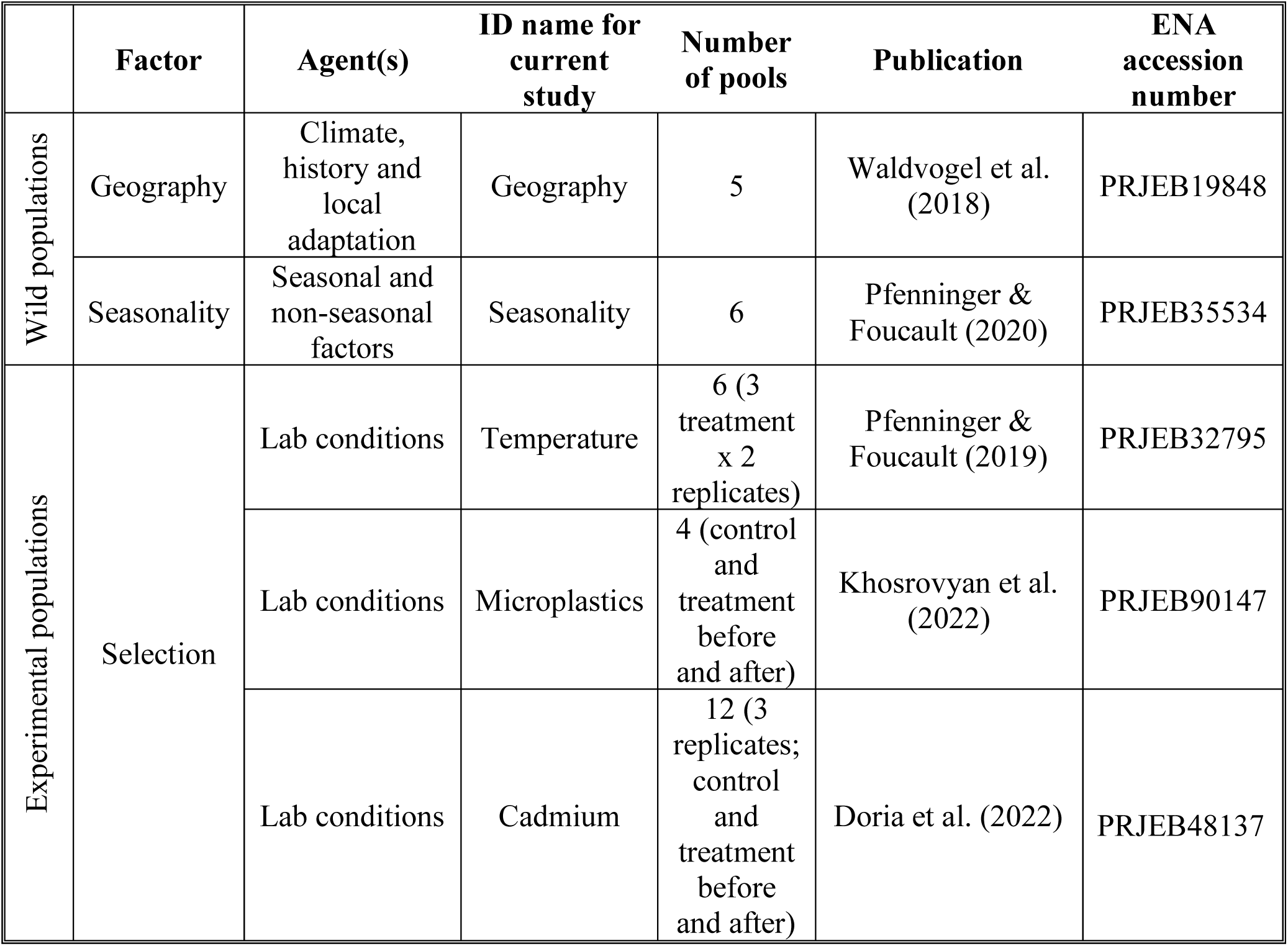
Datasets downloaded from ENA for the estimation of recombination rates.

|  | Factor | Agent(s) | ID name for current study | Number of pools | Publication | ENA accession number |
| --- | --- | --- | --- | --- | --- | --- |
| Wild populations | Geography | Climate, history and local adaptation | Geography | 5 | Waldvogel et al. (2018) | PRJEB19848 |
|  | Seasonality | Seasonal and non-seasonal factors | Seasonality | 6 | Pfenninger & Foucault (2020) | PRJEB35534 |
| Experimental populations | Selection | Lab conditions | Temperature | 6 (3 treatment x 2 replicates) | Pfenninger & Foucault (2019) | PRJEB32795 |
|  |  | Lab conditions | Microplastics | 4 (control and treatment before and after) | Khosrovyan et al. (2022) | PRJEB90147 |
|  |  | Lab conditions | Cadmium | 12 (3 replicates; control and treatment before and after) | Doria et al. (2022) | PRJEB48137 |

**Figure S1.**
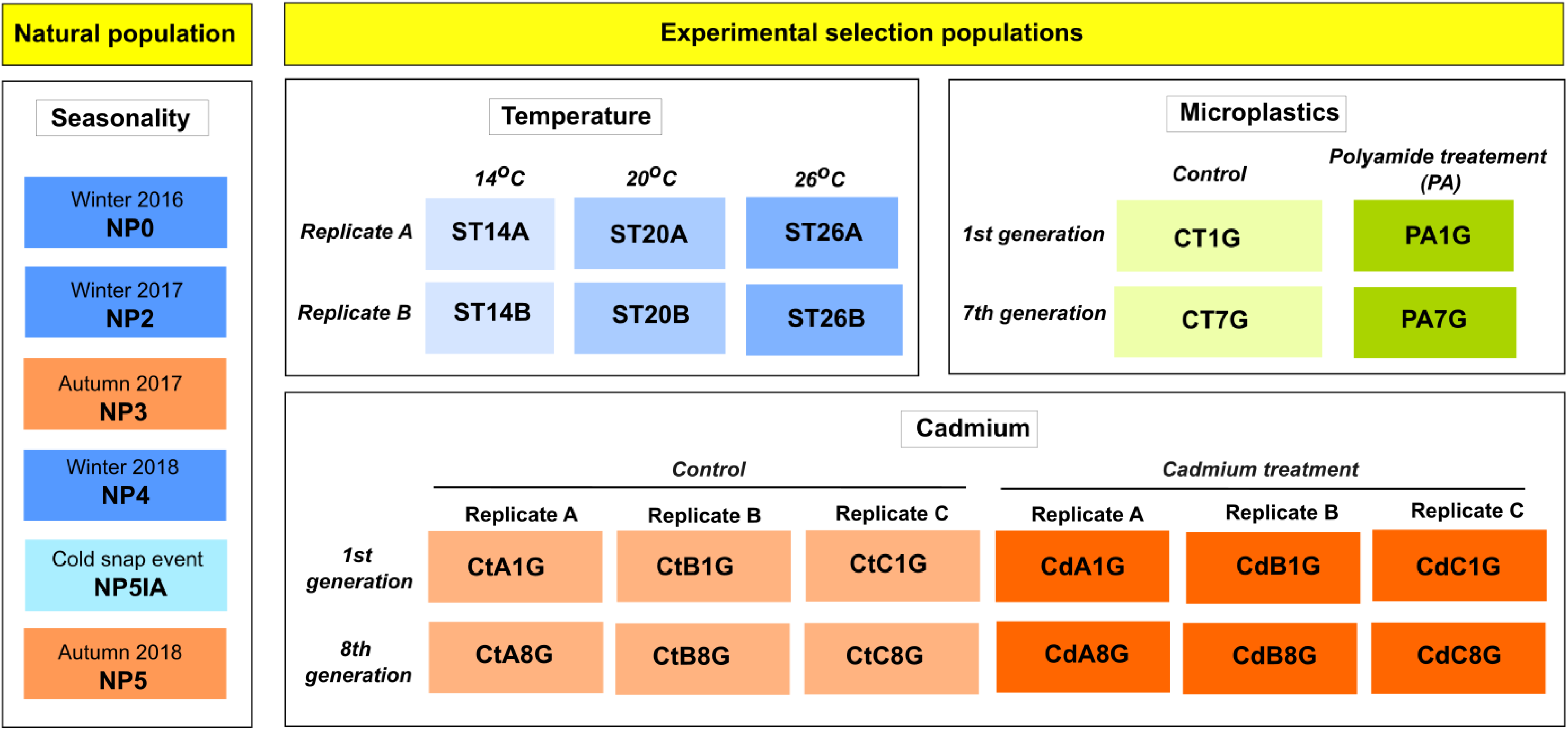
Description of identifiers for pools form seasonality, temperature, microplastics and Cadmium datasets.

**Extraction of GC content.**
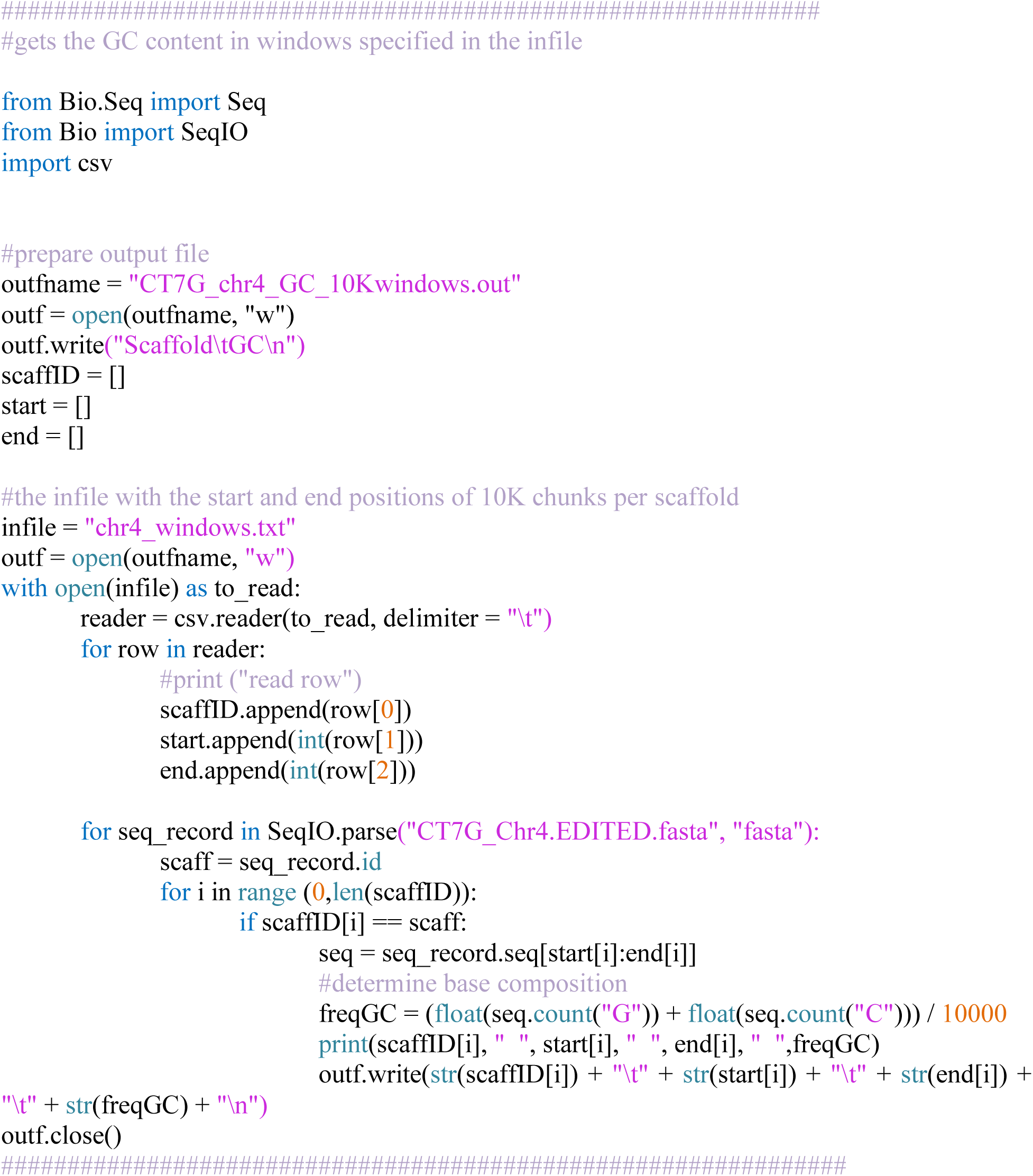
Python script used to extract GC content from each pool using 10kb windows.

## Supplementary Results

**Figure S2.**
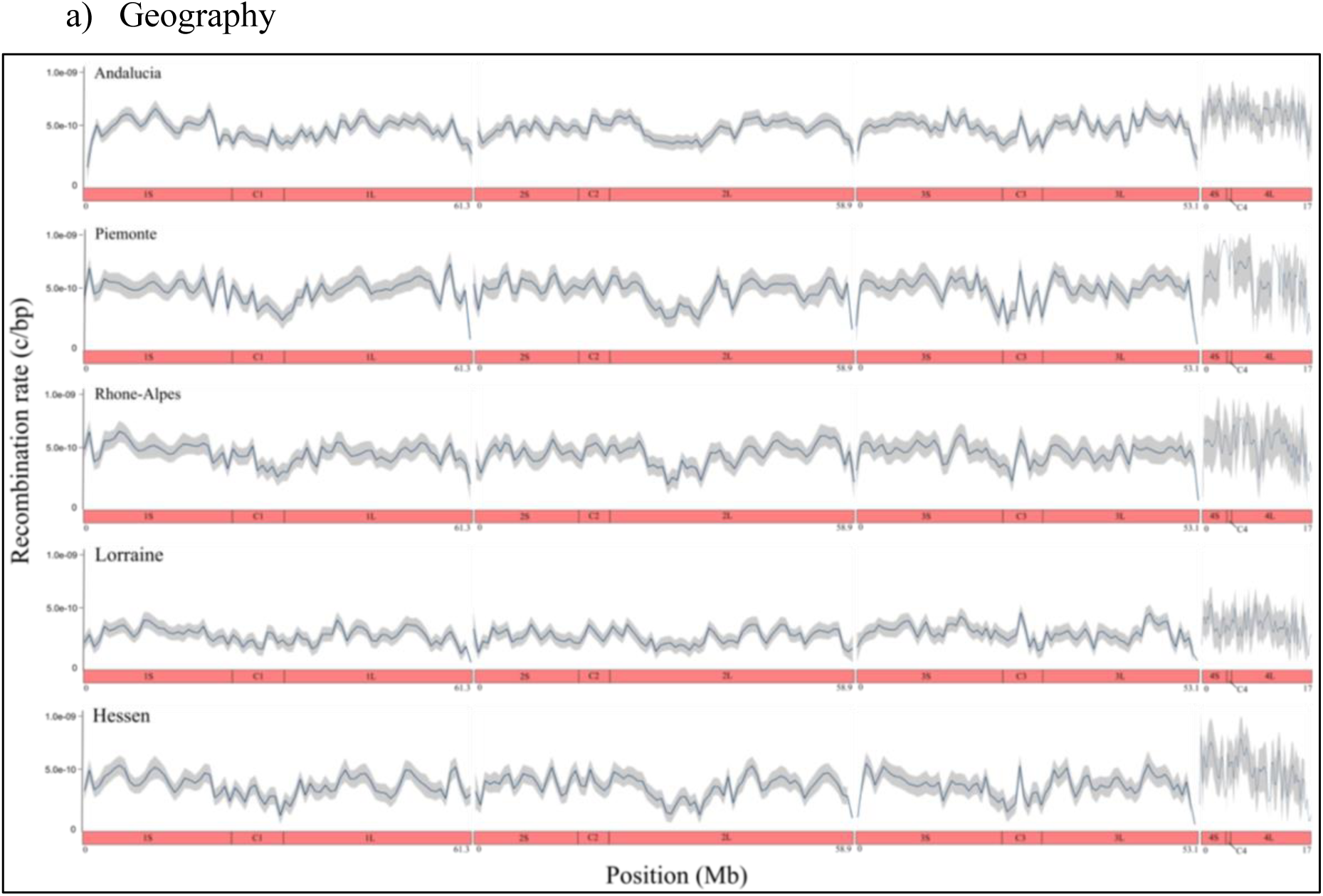
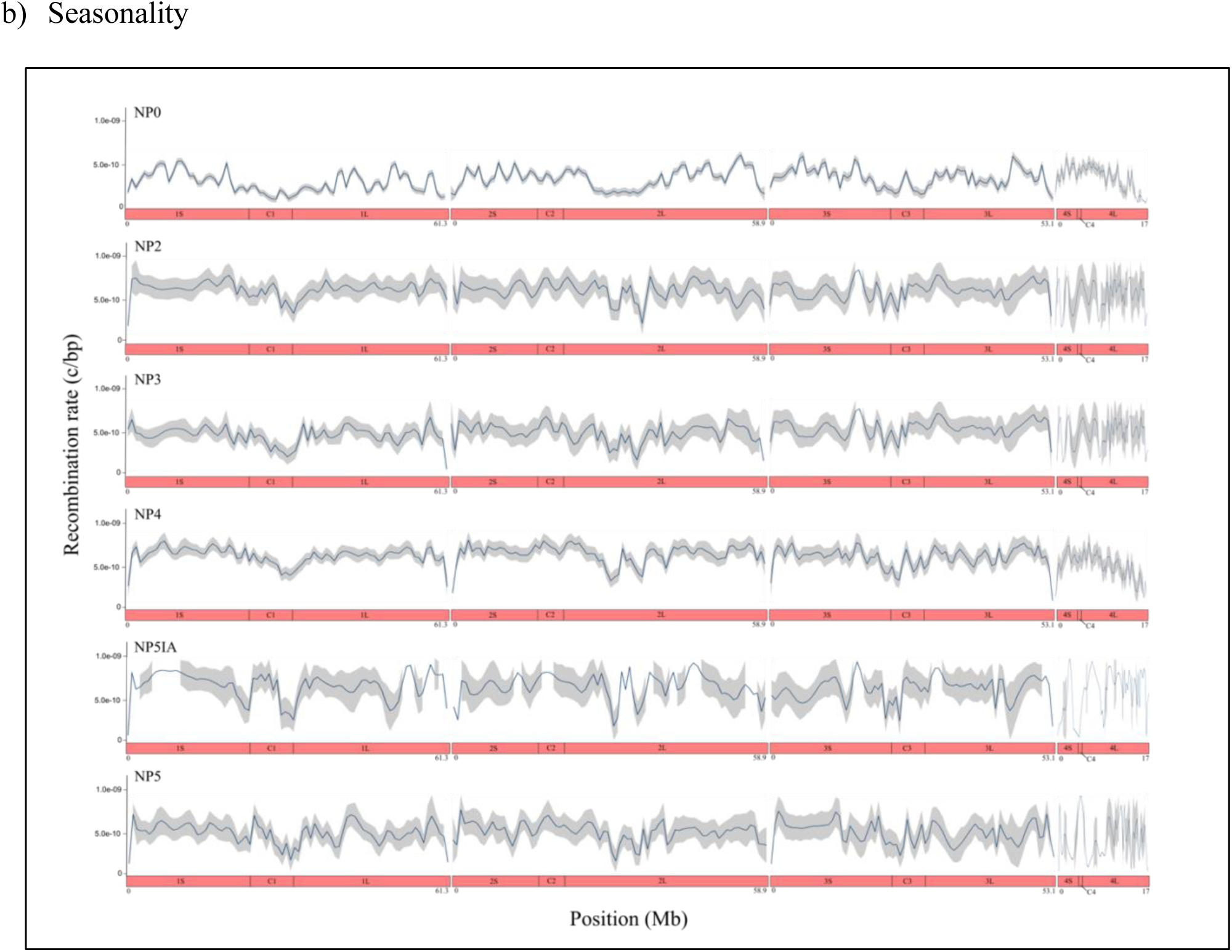
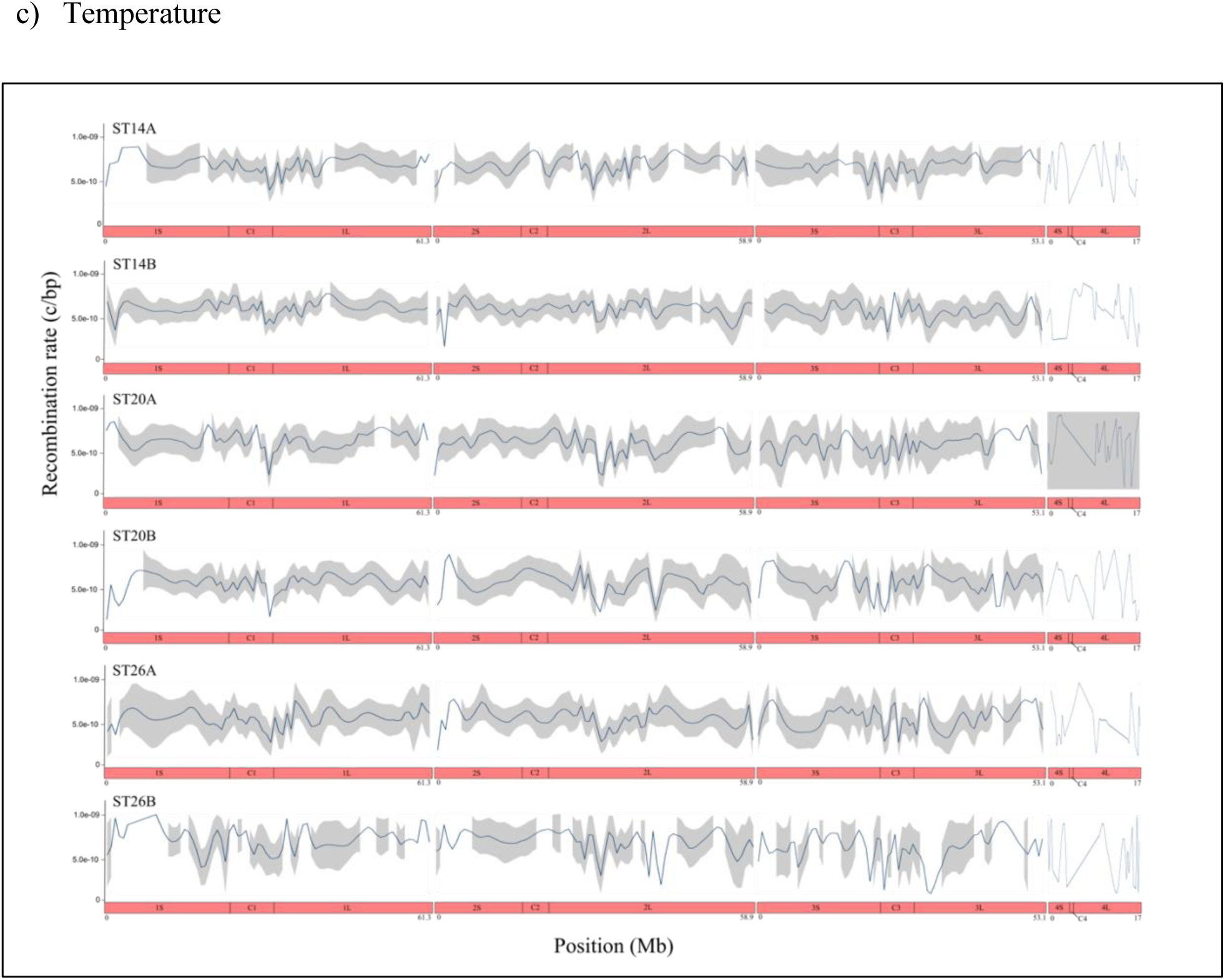
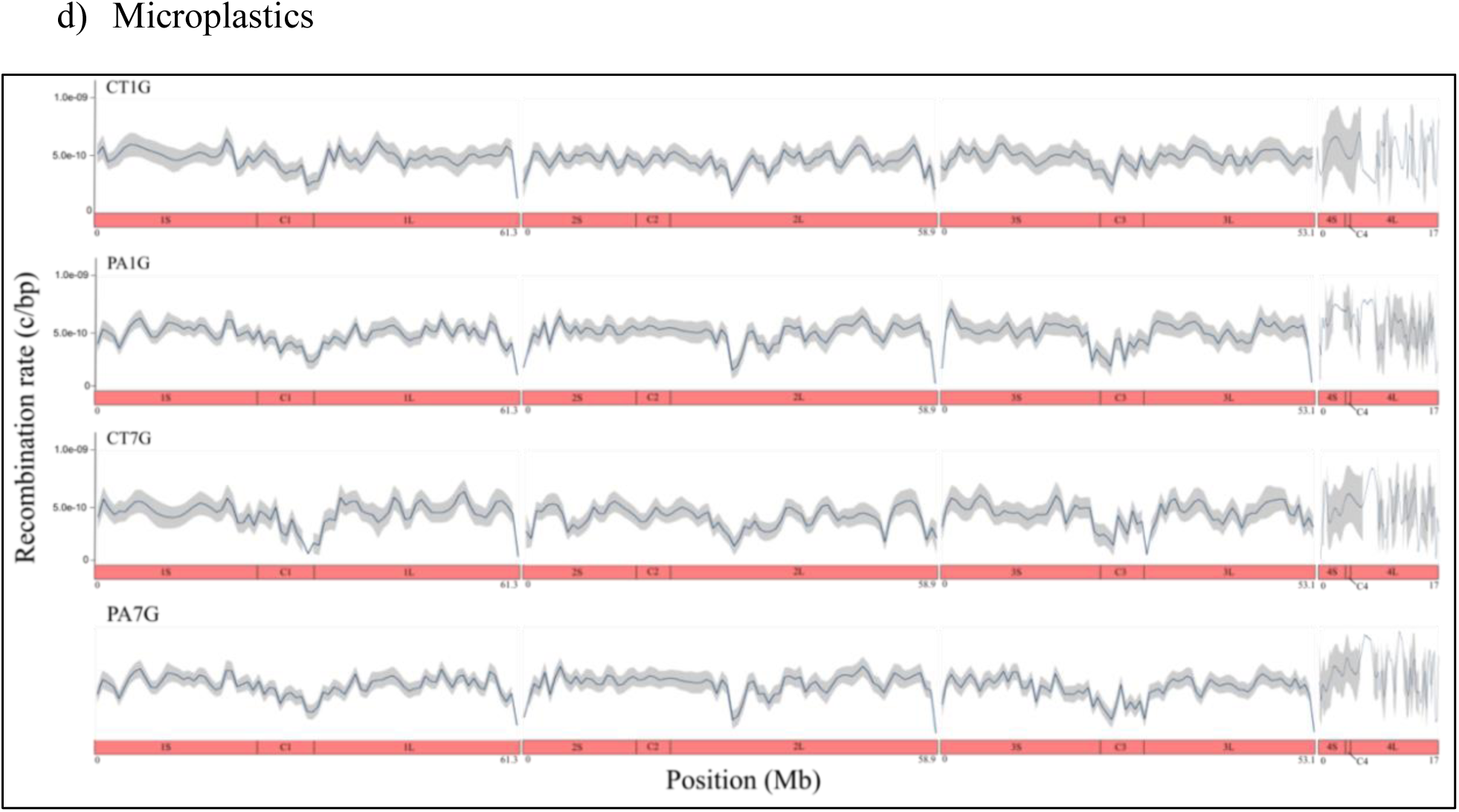
Genome-wide distribution of 10kb recombination rates along four chromosomes from the a) geography, b) seasonality, c) temperature, d) microplastic and e) Cadmium datasets (Mb=million bases).

**Figure S3.**
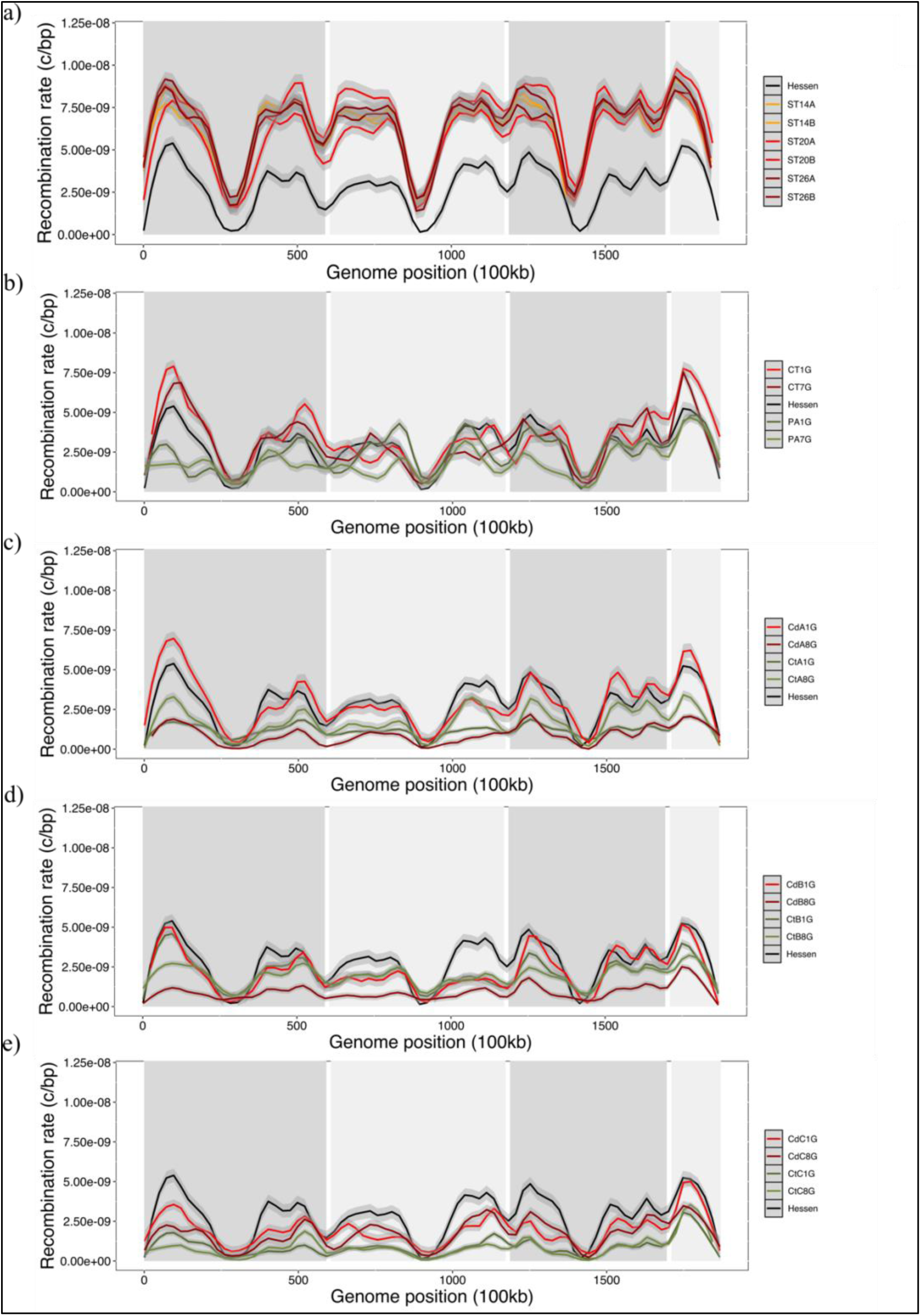
Genome-wide distribution of 10kb recombination rates comparison of Hessen ancestral pool (geography dataset) and experimental selection datasets: a) temperature, b) microplastics and c-e) Cadmium datasets.

**Figure S4.**
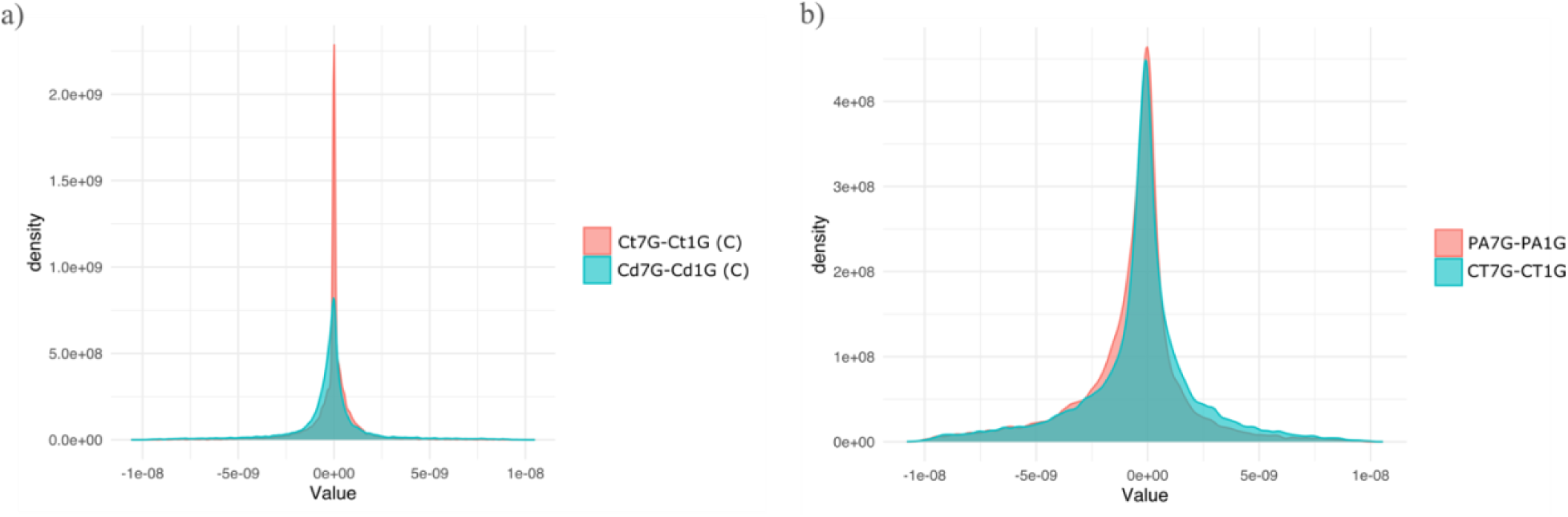
Plots showing the distribution of difference (Δ) in recombination rates between: a) 8^th^ and 1^st^ generation for control and Cadmium pools replicate C; and b) 7^th^ and 1^st^ generation for control and microplastics pools.

**Figure S5.**
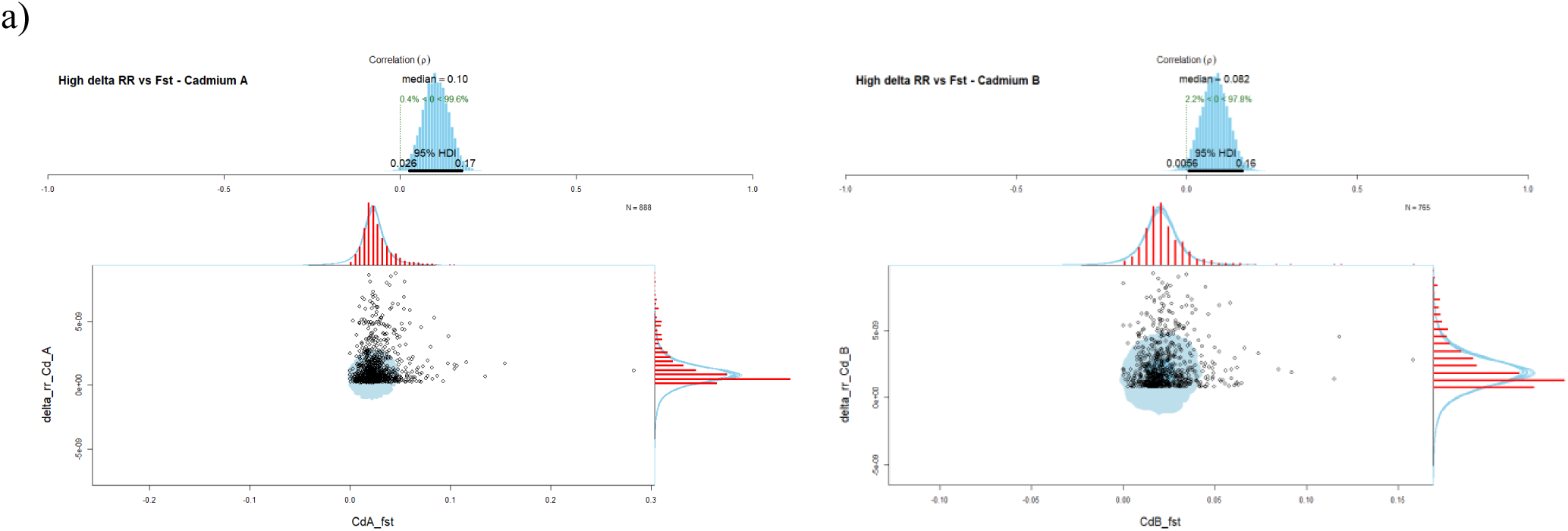
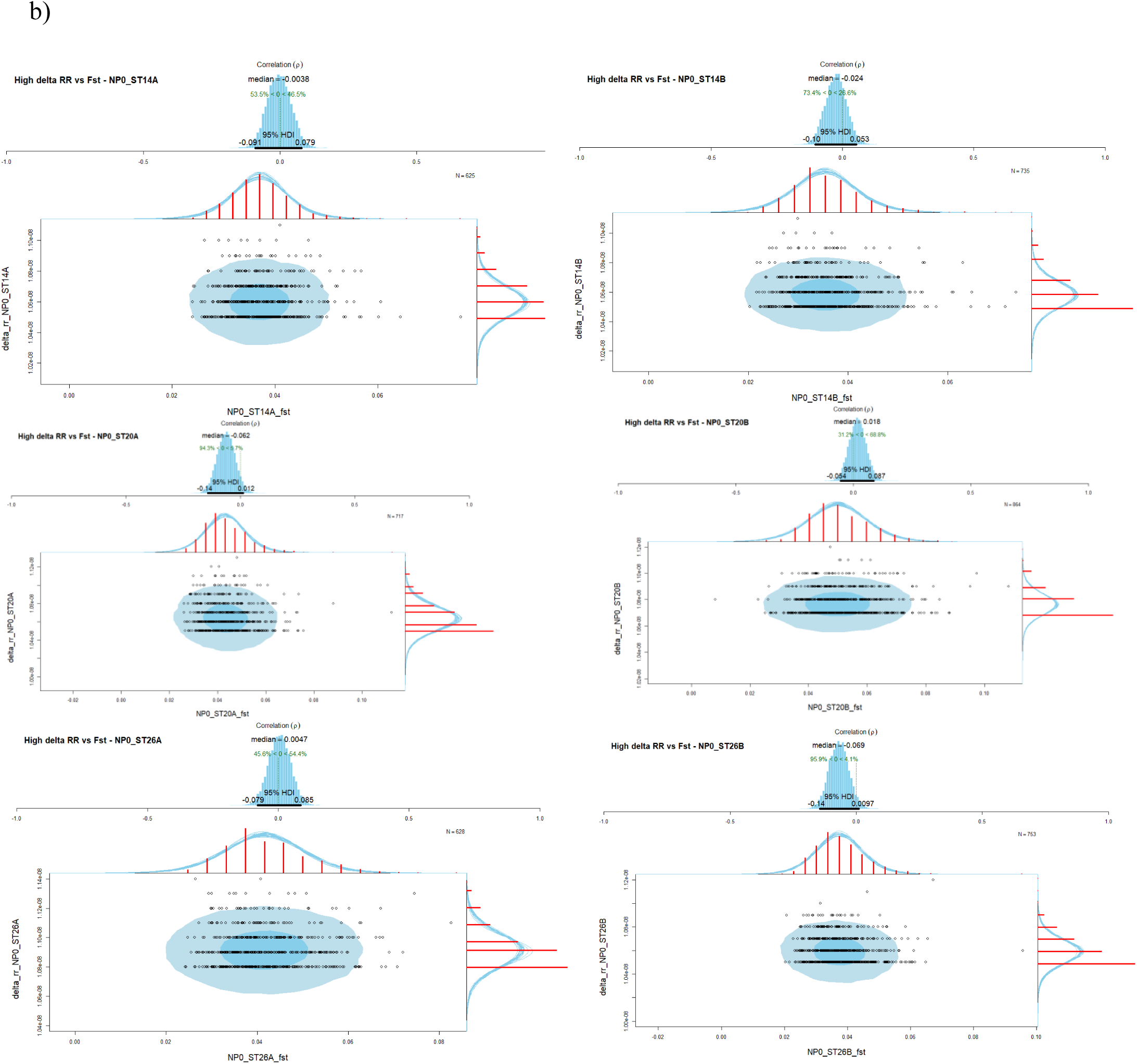
Correlation between high Δ differences of recombination rates (> 95% percentile) and their corresponding F_ST_ window estimate. a) Indicates correlations for Cadmium replicate A and B; and b) indicates correlations all temperature treatments for both replicates and ancestral pool from seasonality dataset (NP0).

**Figure S6.**
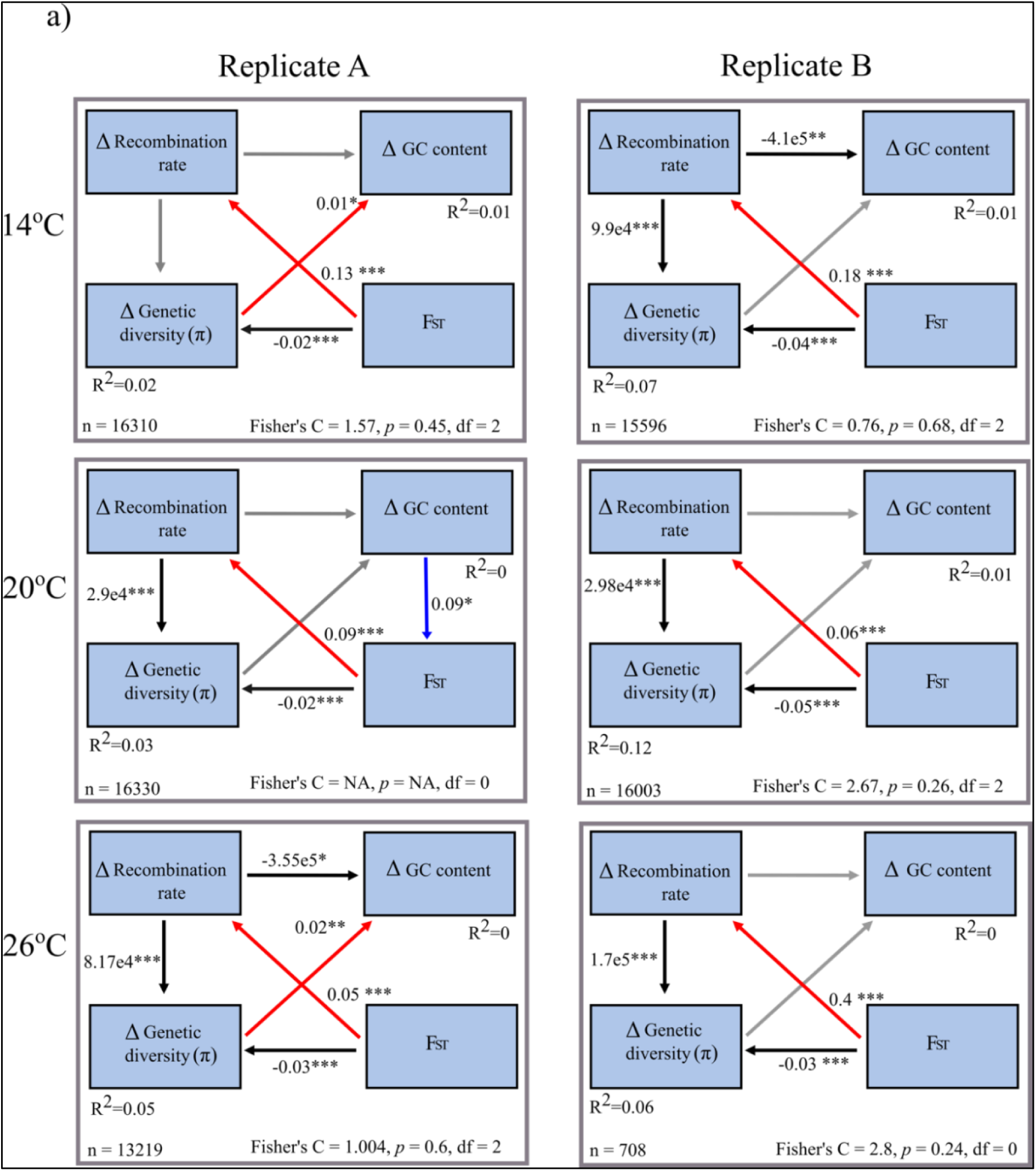
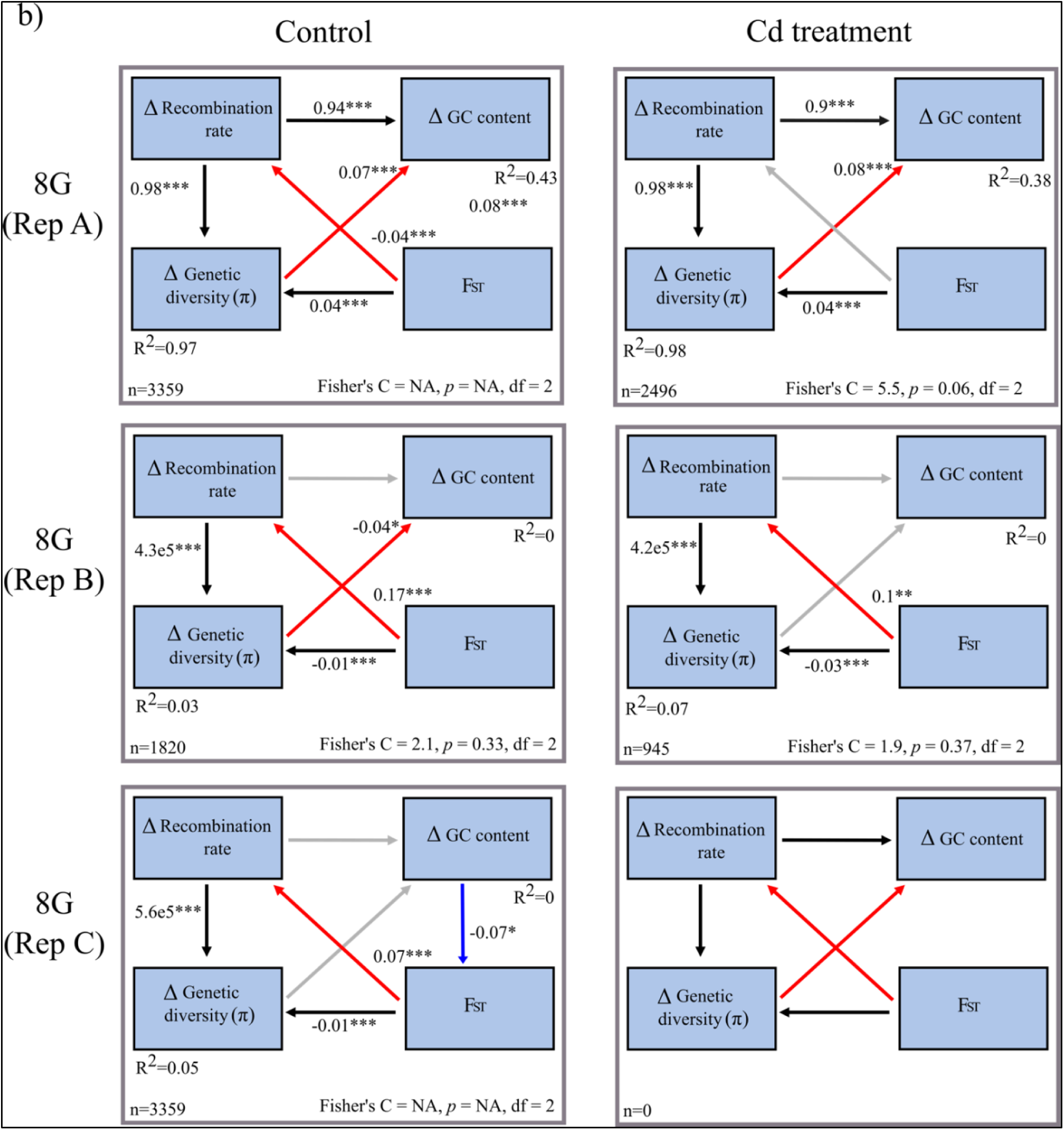
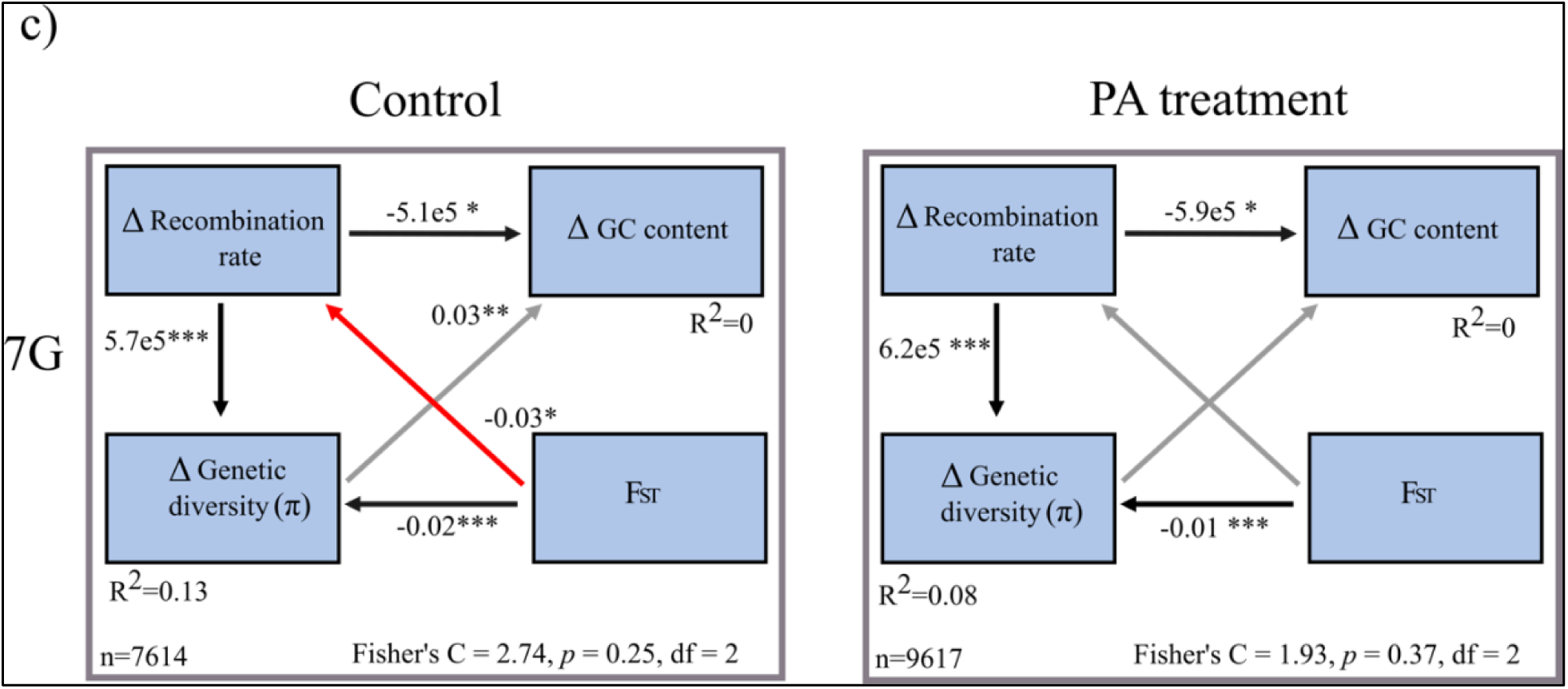
SEM models of hypothesised causal and covariance relationships for experimental datasets: a) temperature dataset; b) Cadmium dataset; and c) microplastics dataset. Fisher’s C indicates model fit, with *p*-values Degrees of freedom are marked with df; n = number of genomic windows included in the specific analysis; Values next to arrows show standardised estimates, with asterisk indicating the statistical significance of the relationship (*p < 0.05, **p < 0.01, ***p < 0.001). Black and red arrows indicate statistical significance causal and covariance relationships, respectively; Grey arrows indicates relationship that were not statistically significance.

**Table S2.** SEM statistical support for causal and covariance relationships based on hypothetical model (Figure 1). Asterisks indicate the statistical significance of the relationship (*p < 0.05, **p < 0.01, ***p < 0.001). ∼ indicates no statistical support. Analysis for Cadmium replicate C 8^th^ generation was not possible dur to missing data. RR = recombination rates, π = genetic diversity, GC = GC content.

|  | Pool | Causal relationship |  |  | Covariance relationship |  |  |
| --- | --- | --- | --- | --- | --- | --- | --- |
| | | RR- $\pi$ | RR-GC | $\pi$ - $F_{ST}$ | RR- $F_{ST}$ | $\pi$ -GC | GC- $F_{ST}$ |
| **Temperature** | 14°C Rep A | ~ | ~ | *** | *** | * |  |
|  | 20°C Rep A | *** | ~ | *** | *** | ~ |  |
|  | 26°C Rep A | *** | * | *** | *** | ** |  |
|  | 14°C Rep B | *** | ** | *** | *** | ~ |  |
|  | 20°C Rep B | *** | ~ | *** | *** | ~ |  |
|  | 26°C Rep B | *** | ~ | *** | *** | ~ |  |
| **Microplastics** | Control 7G | *** | * | *** | * | ~ |  |
|  | MP-PA 7G | *** | * | *** | ~ | ~ |  |
| **Cadmium** | Ct 8G Rep A | *** | *** | *** | ** | *** | *** |
|  | Cd 8G Rep A | *** | *** | *** | ~ | *** |  |
|  | Ct 8G Rep B | *** | ~ | *** | *** | * |  |
|  | Cd 8G Rep B | *** | ~ | *** | ** | ~ |  |
|  | Ct 8G Rep C | *** | ~ | *** | *** | ~ | * |
|  | Cd 8G Rep C | no data | no data | no data | no data | no data |  |

